# COG5-mediated Golgi organization coordinates neural progenitor fate during human brain development

**DOI:** 10.64898/2026.10.27.760061

**Authors:** Martina Polenghi, Elena Restelli, Andrea Jáñez Pedrayes, Bart Ghesquière, Maura Galimberti, Valentina Rava, Giulia Visani, Salma Amin, Cristina Cheroni, Roberta Bosotti, Sara Terzoli, Ikram Taleb Arrada, Olivier Prodhomme, Fanny Alkar, Magdalena Mezaros, Sandrine Vuillaumier Barrot, Arnaud Bruneel, Peter Witters, Marjolaine Willems, Francois Foulquier, Elena Taverna

## Abstract

Apical radial glial cells (aRGs) are highly polarized epithelial cells that drive human cortical development through tightly regulated lineage progression. A distinctive feature of aRGs is their non-canonical Golgi organization, raising the possibility that specialized intracellular trafficking and secretory processes contribute to progenitor behaviour. However, how Golgi architecture contributes to human corticogenesis remains poorly understood.

Here, we investigate this question in the context of COG5 deficiency, a rare congenital disorder of glycosylation associated with neurodevelopmental defects, using patient-derived fibroblasts and brain organoids, N-glycomics, glycolipidomics, and single-cell transcriptomics. We show that COG5 deficiency disrupts Golgi organization and impairs Golgi-to-ER retrograde trafficking. At the neurodevelopmental level, COG5-deficient organoids display altered ventricular organization and reduced basal progenitor abundance. Single-cell transcriptomic analysis reveals rewired developmental trajectories characterized by premature differentiation from apical progenitors with activation of neuronal pathways.

Despite the absence of major abnormalities in plasma glycoproteins glycosylation, glycolipidomic analysis identified marked defects in glycosphingolipid maturation within patient-derived brain organoids, indicating selective impairment of neural lipid glycosylation. Transcriptomic and biochemical analyses further revealed dysregulation of developmental pathways, including reduced Notch signalling and altered localization of both cleaved Notch1 and the Golgi-associated protein ACBD3. Consistent with impaired progenitor maintenance, COG5-deficient organoids displayed altered cleavage plane orientation in apical progenitors.

Together, our findings suggest a link between Golgi organization, intracellular trafficking, membrane glycosylation, and lineage progression in human neural progenitors, identifying Golgi architecture as an important regulator of apical radial glial cell biology and cortical development.

## INTRODUCTION

The formation of the human neocortex depends on the coordinated expansion and differentiation of neural progenitor cells during embryonic development. Among these progenitors, apical radial glial cells (aRGs) represent the primary neural stem cell population of the developing brain and give rise, directly or indirectly, to most cortical neurons (Götz and Huttner 2005; Kriegstein and Alvarez-Buylla 2009). aRGs are highly polarized cells characterized by an apical process contacting the ventricular surface and a long basal process extending towards the basal lamina (Taverna et al. 2016; Noctor et al. 2001). Through tightly regulated symmetric and asymmetric divisions, aRGs balance progenitor maintenance with neuronal production, thereby controlling cortical growth and architecture (Götz and Huttner 2005; Lancaster and Knoblich 2012).The transition from apical progenitors to basal progenitors (the second neural progenitor type to be identified) (Miyata et al. 2004; Noctor et al. 2004; Haubensak et al. 2004), is accompanied by major changes in cell polarity, intracellular organization, and signalling responsiveness (Wilsch-Bräuninger et al. 2012; Polenghi and Taverna 2023; Arai and Taverna 2017). During this process, progenitors progressively lose the apical attachment and acquire distinct morphological and transcriptional identities (Florio and Huttner 2014; Florio et al., 2015.; Kalebic and Huttner 2020). Proper coordination of this transition is essential for cortical development, and disruption of progenitor identity frequently results in neurodevelopmental disorders, with microcephaly as a prominent clinical manifestation (Bizzotto and Francis 2015).

One of the most distinctive features of aRGs is their non-canonical Golgi organization (Taverna et al. 2016). In mammalian cells, the Golgi apparatus typically forms a compact perinuclear ribbon composed of interconnected cisternae (D’Souza et al. 2020). In contrast, in aRGs Golgi elements are distributed along the apical process in discrete units positioned between the nucleus and the apical plasma membrane, aligned with the apico-basal axis of the cell (Taverna et al. 2016). This peculiar organization might support polarized directional trafficking within these highly elongated progenitors, although the functional significance of Golgi architecture in aRG biology remains poorly understood.

Along with its role in protein secretion, the Golgi apparatus represents a central hub for protein and lipids glycosylation, thereby influencing membrane composition, organization and signalling regulation (Agliarulo and Parashuraman 2022). Importantly, Golgi-dependent vesicular trafficking coordinates the spatial distribution and processing of key signalling receptors and membrane components that are essential for developmental decisions (Sheen 2012; Viola et al. 2024; Polenghi and Taverna 2023). Disruptions in the molecular players governing Golgi structure or function have been increasingly linked to neurodevelopmental disorders, suggesting that organization of organelles involved in intracellular trafficking may represent an important layer of regulation during corticogenesis (Rasika et al. 2018).

A major group of disorders associated with Golgi dysfunction is represented by the Congenital Disorders of Glycosylation (CDGs), a heterogeneous class of multisystem diseases caused by defects in protein and lipid glycosylation (Freeze et al. 2015; Jaeken and Matthijs, 2001). Neurological manifestations are among the most common and severe features of CDGs and frequently include developmental delay, intellectual disability, epilepsy, and microcephaly (Paprocka et al. 2021). However, despite the increasing number of identified patients, the relation between Golgi dysfunction and impaired human brain development remains largely unresolved.

The Conserved Oligomeric Golgi (COG) tethering complex is a key regulator of Golgi organization and of vesicular retrograde trafficking (Golgi-to-ER recycling of resident cargo). The complex is composed of eight subunits organized into two lobes, COG1–4 (Lobe A) and COG5–8 (Lobe B), which interact and cooperate in vesicle tethering and membrane recycling within the Golgi apparatus (Blackburn et al. 2019). Defects in individual COG subunits impair intracellular vesicular trafficking compromising both Golgi homeostasis and protein/ lipid glycosylation (Bailey Blackburn et al. 2016; Climer et al. 2015; D’Souza et al. 2020; Reynders et al. 2009; Foulquier 2009). Mutations in several COG subunits have been identified in CDG patients, collectively referred to as COG-CDGs, many of whom present severe neurodevelopmental phenotypes (D’Souza et al. 2020).

COG5 is a core component of lobe B, essential for the stability of other lobe B subunits and, by extension, for the correct orchestration of retrograde vesicular Golgi trafficking. Although previous studies have linked COG5 mutations to developmental abnormalities and microcephaly (Paesold-Burda et al. 2009), the mechanisms by which COG5-dependent Golgi dysfunction affects neural progenitor biology remain unknown.

Here, we investigate the role of COG5 in human cortical development using patient-derived fibroblasts, human brain organoids, N-glycomics, glycolipidomics, and single-cell transcriptomics. We identified a new COG5 mutation associated with neurodevelopmental defects, disrupted Golgi organization, and retrograde trafficking in patient-derived fibroblasts. In patients’ brain organoids, Golgi defects are associated with impaired glycosphingolipid maturation, altered developmental trajectories, and dysregulation of signalling pathways controlling progenitor fate, including Notch signalling.

Together, our findings suggest a link between Golgi organization, intracellular trafficking, membrane glycosylation, and lineage progression in human neural progenitors, identifying Golgi architecture as an essential regulator of NPCs identity and cortical development.

## MATERIALS & METHODS

### Ethics statement

Fibroblasts from COG5-CDG patients and from their unaffected mother were collected by Dr. Willems at Montpellier Hospital and Dr. François Foulquier at the Université of Lille following the patients consent for the collection and usage of biological material for research purposes under local ethical approval. Unrelated control fibroblasts were obtained from the National Facility for Genome Engineering & Disease Modelling, at Human Technopole (Milan, Italy).

#### Fibroblasts culture

Fibroblasts were cultured in DMEM high glucose (ThermoFisher, 11965-092) supplemented with L-Glutamine, 10% FBS (Merck, F2442) and Penicillin (100 U/ml) and Streptomycin (100 μg/ml) (P/S; ThermoFisher, 15140122) and cells were kept in a humidified incubator at 37°C with 5% CO2. Medium was changed once a week. For fibroblasts splitting, the medium was removed, and cells washed once in warm PBS, followed by an incubation with 2mL of Trypsin-EDTA (0.05%), phenol red (ThermoFisher, 25300054) at 37°C for 5 minutes. 10mL of warm conditioned medium were added to the fibroblasts to detach them from the substrate. Cells were then centrifuged for 5 minutes at 300xg. Fibroblasts were resuspended in culture medium and plated on the appropriate support containing maintenance medium as described above. For fibroblasts freezing and thawing, the protocol described for iPSCs was followed (see below) with exception of the culture medium.

#### BFA treatment on fibroblasts

Fibroblasts were plated onto a coverslip. Brefeldin A (BFA) treatment was performed by adding BFA 10nM (Sigma, B6542) to the culture medium and by incubating fibroblasts in an incubator at 37°C with 5% CO2. Fibroblasts were fixed at different timepoints (0min, 5min, 10min, 15min, 20min, 25min and 30min) in 4%(w/v) paraformaldehyde (PFA; Carlo Erba, 387507) for 10 minutes at room temperature (see below) to assess the dynamics of GA fragmentation.

#### Cholera toxin assay on fibroblasts

Fibroblasts were plated onto a coverslip three days prior to performing the experiment. Cells were first placed on ice for 10 minutes and then incubated with 1ug/mL of recombinant Cholera Toxin Subunit B Alexa Fluor™ 555 Conjugate (ThermoFisher) on ice for 30 minutes. Fibroblasts were then washed twice with cold Dulbecco’s phosphate-buffered saline (Euroclone, ECB4004L) without calcium and magnesium (DPBS -/-) and fibroblasts medium was added. Cells were placed at 37°C and then fixed in 4% PFA for 10 minutes at room temperature at the desired timepoints (0 min., 30min., 60 min., 2h, 4h) to track cholera toxin internalization in trafficking organelles.

#### Patients’ fibroblasts reprogramming

Patient fibroblasts were reprogrammed by the National Facility for Genome Engineering & Disease Modelling, Human Technopole (Milan, Italy). Fibroblasts reprogramming into induced pluripotent stem cells (iPSCs) was performed using the Stemgent StemRNA 3rd Generation Reprogramming Kit (REPROCELL, 00-0076), following the manufacturer’s instructions. Briefly, on Day 0, fibroblasts were harvested and seeded at a density of 40,000–50,000 cells per well on Recombinant Laminin iMatrix – 511 pre-coated 6-well plates (amsbio, AMS. 892 012) at a final concentration of 2.4 µg/mL. From Day 1 to Day 3, cells were transfected daily to introduce the reprogramming factors OCT4, SOX2, KLF4, c-MYC, NANOG, and LIN28. From Day 4 to Day 9, cells were cultured in NutriStem™ hPSC XF (REPROCELL, 01-0005) and monitored for the emergence of iPSC colonies. Between Days 9 and 13, primary iPSC colonies were manually picked under phase-contrast microscopy using an EVOS XL imaging system (Thermo Fisher Scientific). A total of 60 colonies per sample were picked and plated into 96-well plates (Passage 0). Starting from passage 1, the clones were transferred into 24-well plates and maintained for 10 passages. At passage 10, the 15 best-performing clones—selected based on proliferation rate and morphology—were cryopreserved.

Among these, up to 6 clones were selected for genome integrity analysis to confirm the absence of chromosomal aberrations introduced during reprogramming. This analysis was performed using low-pass whole-genome sequencing. The 3 best-performing clones were then subjected to a full panel of quality control assays, including: expression of pluripotency markers, differentiation potential into the three germ layers, STR analysis for cell line authentication verifying the match with parental cell line, genome integrity by low-pass whole genome sequencing to exclude large aneuploidies or chromosomal abnormalities, mycoplasma testing, and post-thaw recovery assessment after amplification for masterbank cryopreservation. The resulting QC documents for all the clones used in this work is shown in Annex I. For each of the 3 validated clones, a master cell bank consisting of 10 cryovials was generated and delivered to the end-user of the facility.

#### iPSCs maintenance

Human brain organoids were obtained from WTC11 (control iPSCs; Coriell), HMGU1 (control iPSCs; Helmholtz Institute, Munchen), and patients’ iPSCs. Female and male COG5 patients have been labelled for this work as COG52 and COG53, respectively. The iPSCs were cultured on 6-well plates coated with Recombinant Laminin iMatrix - 511 and kept in mTeSR1 medium (StemCell Technology, 85850) in the incubator at 37°C and 5% CO2, with daily medium change. The cells were passaged once a week, when 80% confluency was reached, by washing once with DPBS -/-, and incubation with ReLeSR (StemCell Technology, 100-0484) for 30 seconds at room temperature. ReLeSR was removed and cells were placed in the incubator for 5 minutes. A 1ml of fresh mTeSR1 was added and used to detach cells through two to three pipetting to maintain intact colonies. The split volume of cell suspension was transferred to a new well, in a final volume of 2ml of mTeSR1 with iMatrix to allow cell attachment.

#### Thawing iPSCs

To thaw iPSCs, the cryogenic vial from liquid nitrogen tank was quickly put in the water bath at 37°C for 1 minute. Cells were transferred, with a wet pipet tip, into a tube already containing 5ml of mTeSR1. The cell suspension was centrifuged for 5 minutes at 300xg. After centrifugation, the supernatant was discarded, and the cell pellet was resuspended into 1ml of mTeSR1 with ROCK inhibitor Y-27632 (RI; Tebu-bio/Peprotech, 1293823). Then, 1/3 and 2/3 of cells volume were plated into new 6-cm wells plates with a final volume of 2ml mTeSR1 supplemented with RI for cell attachment.

#### Freezing iPSCs

Cell medium was removed, cells were washed with DPBS -/-, and incubated for 5 minutes at 37°C with 1ml of ReLeSR. Cells were detached from the plate with 2 ml of mTeSR1 and pipetted two to four times. The cell suspension was transferred to a tube and centrifuged for 5 minutes at 200xg. The supernatant was discarded, and the cell pellet was resuspended with 500µl of mFreSR (StemCell, 5854), transferred to a cryogenic vial and progressively frozen to -80°C in a freezing box. After 24h cells were transferred in liquid nitrogen for long term storage.

#### Human brain organoids culture

Human brain organoids were generated following Lancaster protocol (Lancaster and Knoblich 2014) with minor modifications on cell concentration and time windows (Supplementary Fig. 4). Briefly, iPSC at 70-80% of confluency were washed once in DPBS -/-, detached with Accutase (Sigma, A6964) for 5 minutes at 37°C, and diluted in 2 mL of mTeSR1. Cells were centrifuged for 5 minutes at 200xg and resuspended in 1mL of mTeSR1 supplemented with RI. Cells were counted and 9x10^3^ cells/well of a U-bottom 96 well plate (Twin Helix, MS-9096UZ) were added to mTeSR1 with RI medium. Cells were then seeded adding 200µL of suspension to each well of the 96 well plate to form Embryoid bodies (EBs). Media change was done every other day using mTeSR1 without RI. At day 4, medium was changed to Neural induction medium (NIM was prepared with DMEM F12 (ThermoFisher, 31330038) and Neurobasal (ThermoFisher, 21103049) in ratio 1:1, N2 1:200 (ThermoFisher, 17502048), Glutamax 1:100 (ThermoFisher, 35050038), 1µg/mL of Heparin (Sigma, H3149-25KU), 0.5% of MEM NEAA (ThermoFisher, M7145-100ML) and P/S 1:100 with a volume of 150µL per well. Media change was done every other day. At day 7 the EBs were embedded in Geltrex (ThermoFisher, A1413302) droplets (one drop of 50 ul per EB) and put in 6cm petri dishes in Differentiation medium without Vitamin A (DM-VitA was prepared with DMEM F12 and Neurobasal in ratio 1:1, B27 -VitA 1:100 (ThermoFisher, 12587010), N2 1:200, Glutamax 1:100, 2.5µg/mL Insulin (Sigma, I9278), 50uM 2-mercaptoethanol (Merck, 8057400005) and 0.5% of MEM NEAA and P/S 1:100. Media change was done every other day until day 13 when the embedded EBs were transferred in DM+VitA (same composition of DM-VitA but with B27+VitA 1:100 (ThermoFisher, 17504044)). Organoids were then kept in culture until the desired stage by placing the petri dish on a shaker set at 70rpm in the incubator.

#### Organoids’ electroporation

We performed electroporation on human brain organoids to reconstitute intracellular architecture and organelles position in single GFP-labeled cells.Organoids at d40 were collected in DM+VitA without P/S prior to electroporation. The solution containing 1µg/µL pCAGGS-GFP plasmid in H2O was kept on ice until use. Borosilicate glass capillaries (O.D. 1.2mm, I.D. 0.94mm. 10cm length, Sutter Instruments) were pulled using a needle Puller (Model P-1000 Sutter Instrument) with the following set up: Heat 500, Pull 60, Vel 80, Delay 100 and Pressure 280. Organoids were moved to PBS and ventricles were identified by placing the organoids under a dissectoscope equipped with magnifying lenses. The plasmid was injected into single ventricles using the glass capillary under visual inspection. For the electroporation 5 pulses of currents were applied at 80V for 50ms each at 500ms intervals using an electroporator (Ecm830, BTX). Electroporated organoids were moved to the differentiation medium without P/S for 4h and then placed back in normal culture medium. Electroporated organoids were kept in culture for 48h to allow for the expression of the reporter gene and the subsequent reconstruction of the Golgi apparatus in apical progenitors.

#### Cryosections

For cryosectioning, organoids were fixed with 4% PFA for 40 minutes, washed in DPBS (-/-) and infiltrated with 30% sucrose (Sigma, S0389) at 4°C overnight prior to embedding in Killik O.C.T. compound (Bio optica, OCT 05-9801) in embedding molds (Fisher Scientific, 50-465-348). Embedded organoids were then frozen in n-pentane (Carlo Erba, 468121) and stored at −80 °C. 20µm-thick sections were cut using a cryostat (Leica Biosystems, LEICA CM1950) and resulting sections were stored at -20°C for further processing.

#### Immunofluorescence staining on fibroblasts and human brain organoids

##### Immunofluorescence staining on human fibroblasts

For immunofluorescence staining on fibroblasts, permeabilization with Triton-X-100 0.05% (Merck, T8787) in PBS for 10 minutes was done, followed by 1h incubation with blocking solution containing 0.2% Gelatin (Roth, 4308.1), 0.05% Triton X100/PBS, 30 mM NaCl at RT. Cells were then added with the Ab I in blocking solution and kept o/n at 4°C with gentle shaking overnight. The next day cells were washed 3 times for 5 minutes each in PBS and the secondary antibodies were added in blocking solution with 1ug/ml DAPI (ThermoFisher, 62248) as nuclear counterstain for 1h at RT. Cells were washed in PBS and, if on coverslips, the sample was mounted with Mowiol 4-88 (Sigma, 81381).

##### Immunofluorescence staining human brain organoids

Cryosections were dried for 5 min at RT and then permeabilized with 0.5% Triton-X100 in PBS for 40 min. Subsequently, the procedure was followed as for fibroblasts with more and longer washes in PBS and using as blocking solution 0.2% Gelatin, 0.3% Triton X100/PBS, 300 mM NaCl. Stained sections were mounted in Mowiol and imaged.

A complete list of all primary, secondary antibodies and counterstains used, along with the corresponding concentrations are listed in the antibodies section in Table 1.

**TABLE I.**
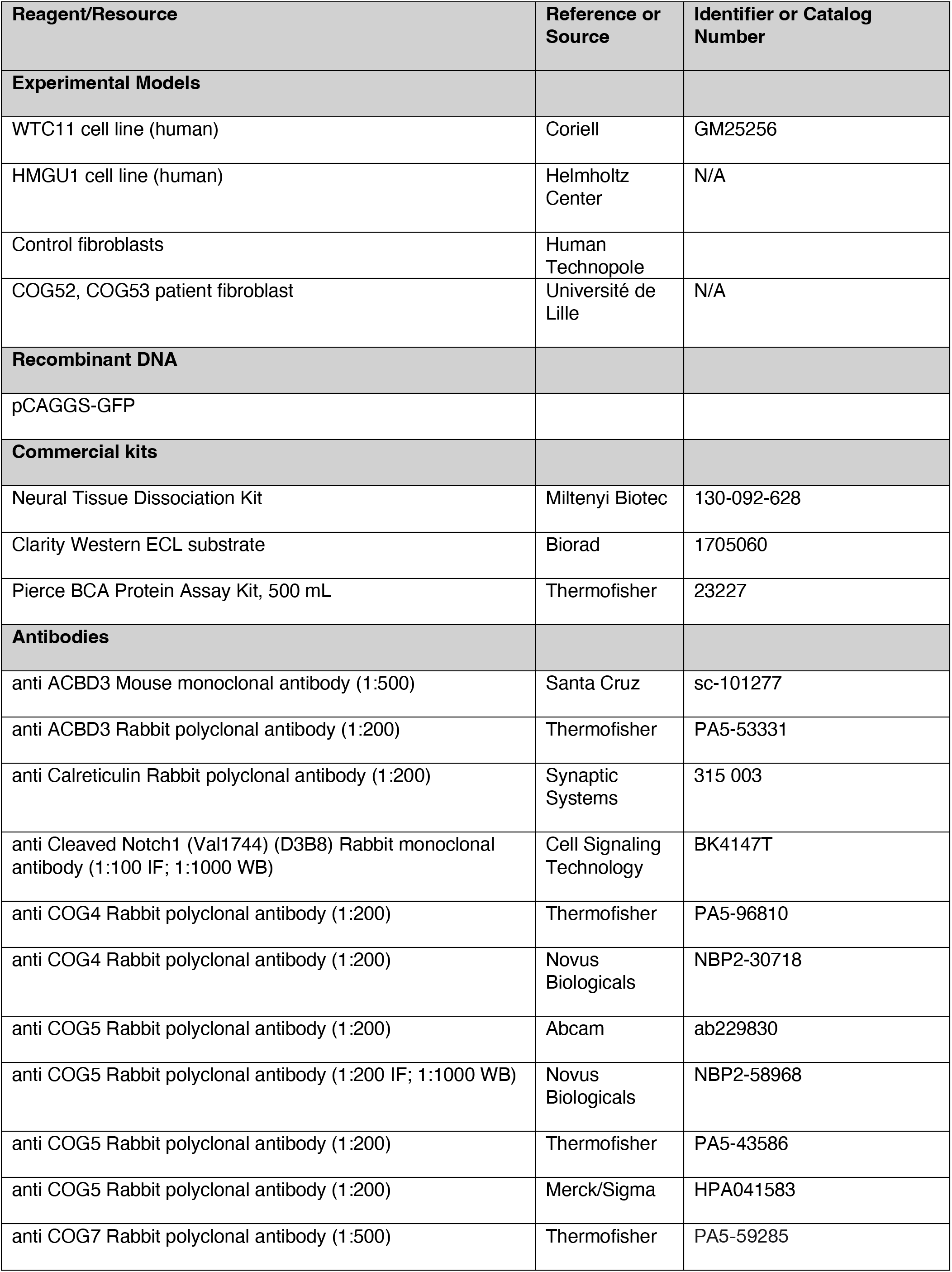
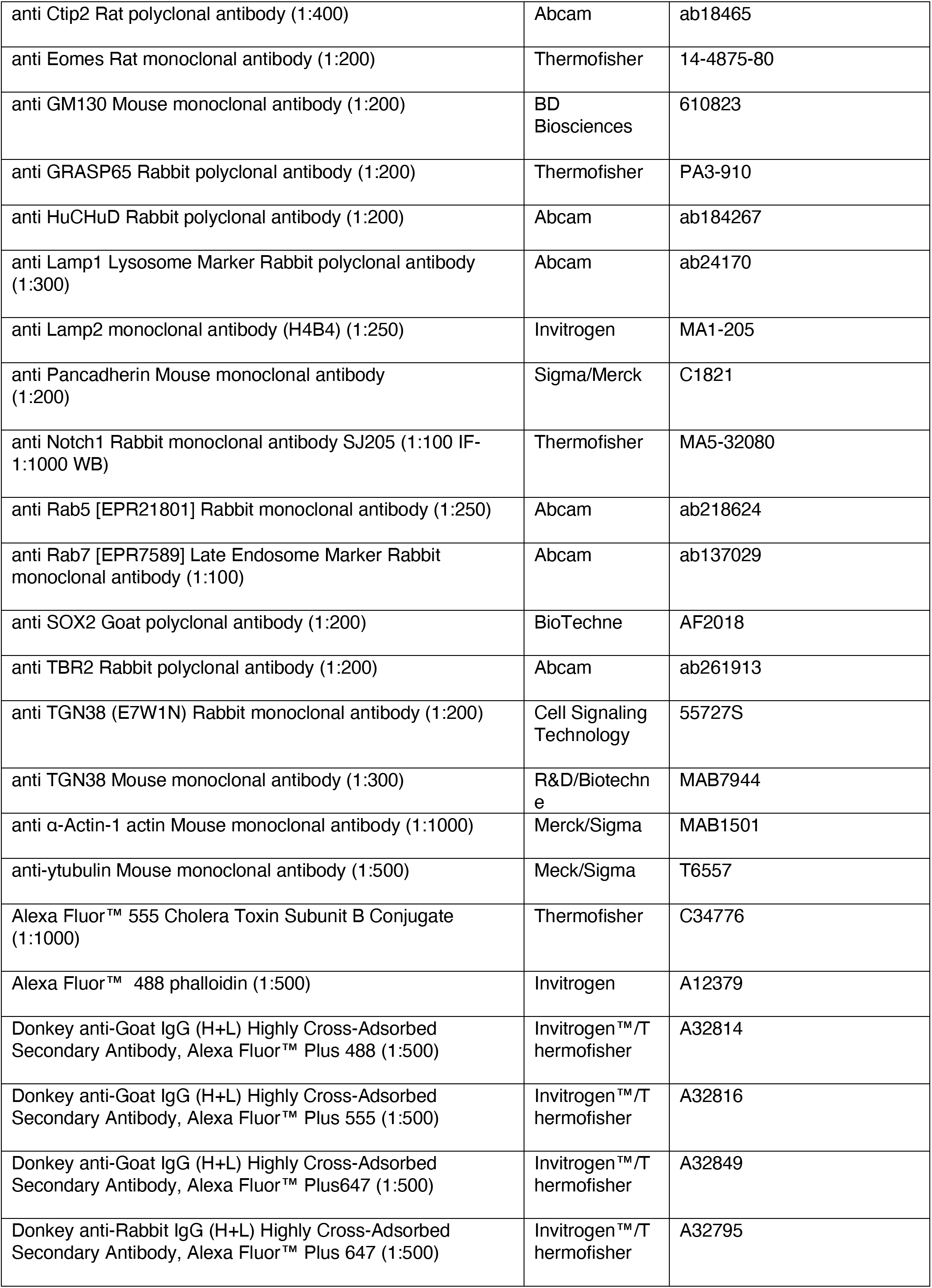
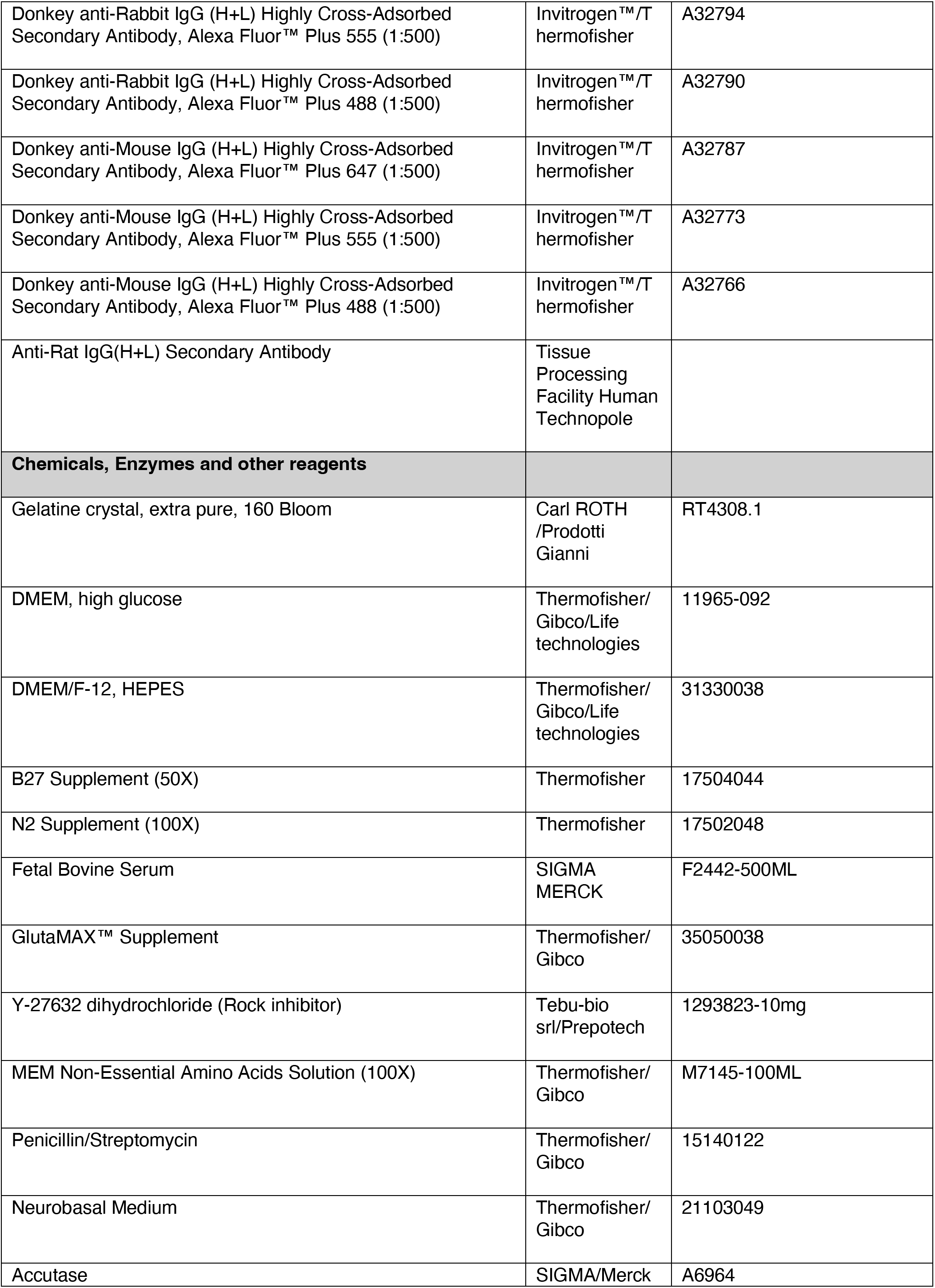
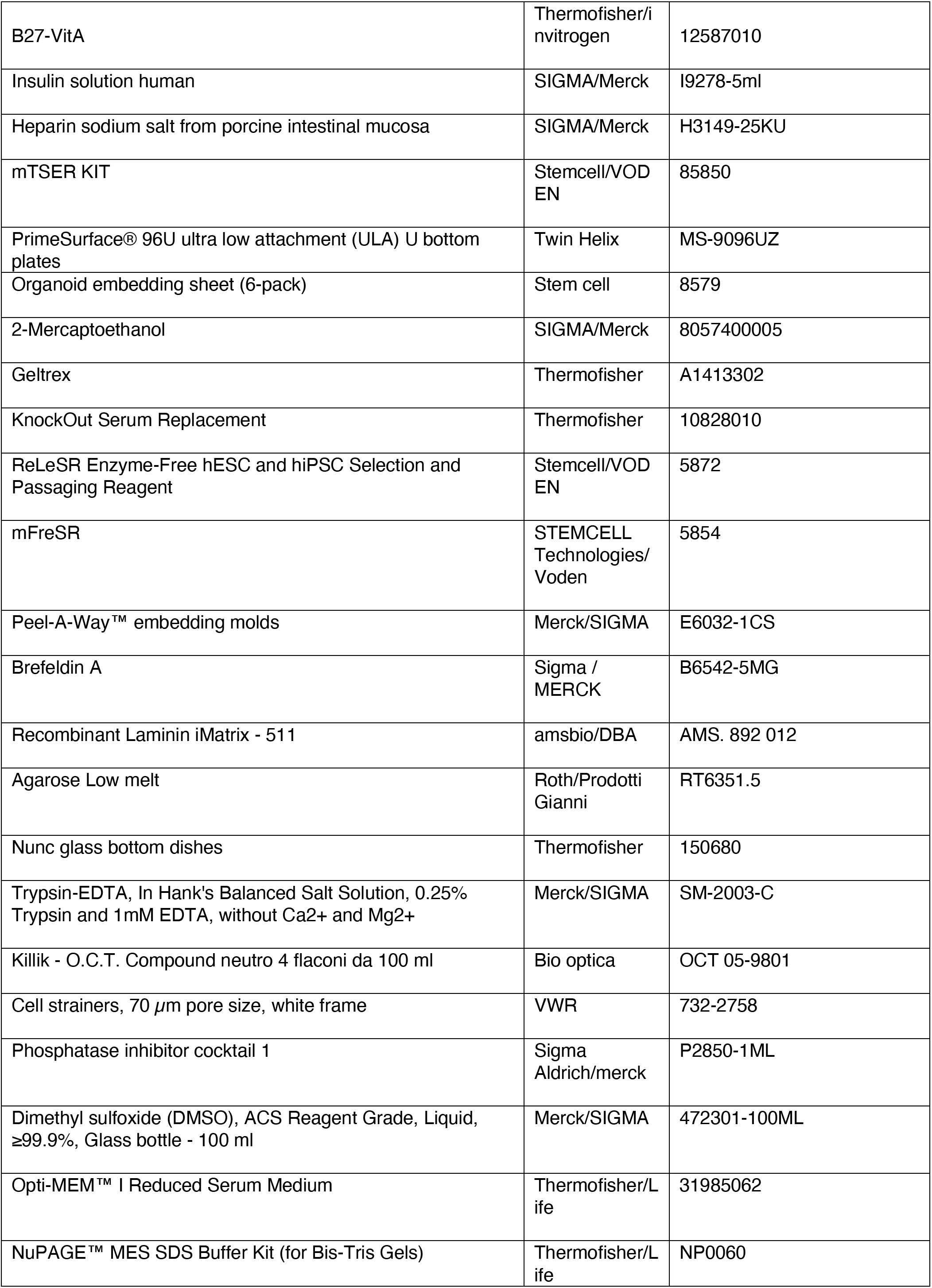
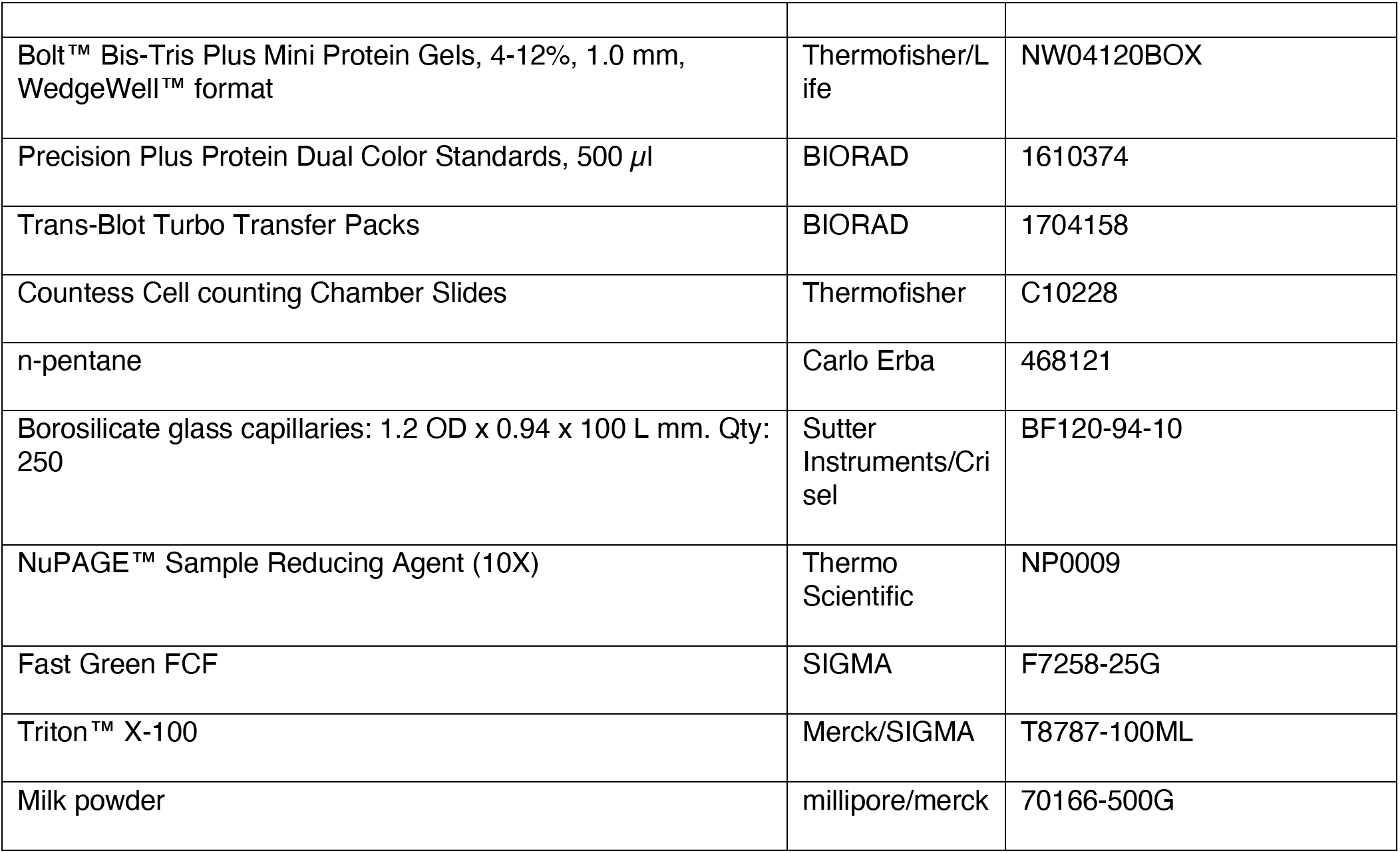

#### Image acquisition

Imaging was performed using a widefield fluorescence microscope (Axio Vert A1, Zeiss, Germany), a confocal microscope (Zeiss LSM 780 NLO; Zeiss, Germany) or a spinning disk confocal microscope (CrestOptics). Confocal images shown here are 0.5 µm-thick single optical sections, unless stated otherwise. Confocal images were obtained using either a PlanApo 63X oil or PlanApo40X oil immersion objectives. 405nm, 488nm, 561nm and 639nm laser lines were used. All confocal images shown here are 0.5 µm-thick single optical sections, unless stated otherwise. Samples were also acquired using a spinning disk confocal microscope, that comprised a Nikon Ti2-E inverted microscope equipped with a CrestOptics V3 Light, which was coupled to Photometrics Prime 95B 25 mm camera, configured with a 50-micron pinhole, and to a Celesta Lumencor as lightsource. The imaging process was conducted using a PlanApo LambdaD FN25 DIC 10X/0.45 and PlanApo LambdaS 40X/1.25 Sil objective, capturing Z-stacks with a 0.5-micron Z-step size to encompass the complete volume of the sample. 405nm, 488nm, 561nm and 639nm laser lines were used.

#### Image analysis

**I**mmunofluorescence images of single optical sections were quantified using Fiji (ImageJ2 v. 2.16.0/1.54p) (Schindelin et al. 2012)

##### Fate markers quantification

Cell quantifications for nuclear fate markers were performed in Fiji either manually with the CellCounter function or using StarDist 2D plugin considering all the area corresponding to an organoid ventricle. The percentage of cells was calculated by dividing the number of cells positive for a given nuclear fate marker for the total number of DAPI-positive nuclei/cells.

##### Organelles quantification

Reconstruction of subcellular features in single cells was performed by manually segmenting the cell contour and cell boundaries (as assessed by the GFP expression) and the organelle of interest (as highlighted by the specific organelle’s marker signal) on the single optical planes from the full Z-stacks covering the entire cell’s volume. Cell reconstruction was then performed, and organelles area and organelles’ number were measured by annotating and analyzing the corresponding ROI (region of interest). When calculating the distance from the apical surface, a segment perpendicular to the apical surface was drawn and its length measured.

##### Colocalization quantification

JaCoP plugin was used to quantify the colocalization of two given channels. This tool was specifically used to quantify (i) the colocalization of ACBD3 with trafficking pathway organelles and (ii) the colocalization of CTB with Golgi (iii) the colocalization of Notch and NCID with Golgi and lysosomes.

##### Mitotic spindle orientation

Mitotic spindle orientation in dividing APs was determined by measuring the angle between the plane of division (as assessed by DAPI signal) and the ventricular surface (identified by Phalloidin or y tubulin) using the angle tool in Fiji. The percentage of vertical (60°-90°), oblique (30°-60°) and horizontal (0°-30°) cell divisions was calculated and compared between the WT and the other conditions. Final image panels were processed and assembled using Adobe Illustrator (v. 30.4)

#### scRNAseq

For scRNAseq, the Chromium GEM-X technology (10x Genomics) was used following published protocols from Illumina. For sample preparation d40 organoids from WTC11, COG52 and COG53 lines were used. 3 independent differentiations experiments with pools of 8 organoids from each experiment and line were used for scRNAseq. Upon pooling of organoids, the tissue was dissociated using the neural tissue dissociation kit (Miltenyi Biotec, 130-092-628). Briefly, 200µl of enzyme P resuspended in buffer X was added to the organoids that were then incubated at 37°C in waterbath for 15 to 30 minutes. After incubation 6µL of enzyme A were added to stop the reaction and full organoids dissociation was achieved through manually pipetting up to 10 times. The suspension was then passed through a 70µm filters (VWR, 732-2758) and the collected cells were briefly centrifuged at 300g for 2 minutes and finally resuspended in 5% FBS (Merck, F2442) in PBS before being handled to the National facility for Genomics (Human Technopole) for further steps. Cell viability was assessed as quality control and only samples with >98% viability were processed. Briefly, single cells were encapsulated into Gel Emulsion Droplets using the Chromium X instrument (10x Genomics) targeting 5’000 cells for each sample. To optimize costs and efficiency, three or four samples were multiplexed in a well of the 10x chip following the instructions of the GEM-X Universal 3’ Gene Expression v4 4-plex Protocol (CG000768, 10x Genomics). From each pool, 3’ Gene Expression libraries were generated in accordance with the 10x Genomics Protocols. Libraries were pooled at a ratio dependent on the 10x Genomics indication and loaded on the Novaseq6000 sequencer to achieve a minimum of 50’000 reads/cell for each library.

#### scRNAseq data analysis

##### Differential gene expression (DGE) analysis

For the differential expression analysis, a pseudobulk approach was applied, in which raw transcript counts were summed across all cells belonging to each combination of biological sample (*sample_id*) and cell cluster (previously defined by Leiden clustering). This strategy allowed the aggregation of technical replicates, thereby reducing spurious variability. Based on the cell type annotation performed previously, the clusters of interest were selected: cluster AP (Apical progenitors), cluster IPC_N (basal progenitors), and clusters N_1, N_2, Imm_N (neurons). Differential gene expression (DGE) analysis was then performed using the DESeq2 package (v1.48.2) in R. Only genes expressed in at least 10% of the samples were retained. A DESeq2 object was defined, and the model was fitted using the function DESeq(test = "Wald", fitType = "parametric"). Results were extracted for the contrasts of interest (COG52+COG53 vs WT, COG53 vs WT, COG52 vs WT). Significantly regulated genes were identified based on both statistical significance (*padj* < 0.05) and fold-change threshold (log2FoldChange| > 1.5). Up- and down-regulated genes were then visualized through volcano plots generated with the *ggplot2* package (v4.0.0) in R, highlighting genes with log2FoldChange > 1.5 & padj < 0.05 as upregulated and those with log2FoldChange < -1.5 & padj < 0.05 as downregulated.

##### Differential abundance (DA) analysis

Differential abundance (DA) analysis was performed using Milopy, a Python implementation of the Milo framework (v0.1.1). The KNN graph was already constructed in the Scanpy workflow using Harmony-corrected principal components (sc.pp.neighbors(mdata["rna"], use_rep="X_pca_harmony", k=100, n_pcs=12)), where k indicates the minimum neighborhood size for DA quantification. Neighborhoods were defined by selecting 10% of all cells as centers (milo.make_nhoods(mdata["rna"], prop=0.1)), grouping cells into overlapping local regions on the graph. The number of cells in each neighborhood per sample was counted (milo.count_nhoods(mdata, sample_col="sample_id")) and used for differential abundance testing with a generalized linear model. Comparisons included COG52 vs WT, COG53 vs WT, and COG52+COG53 vs WT. Neighborhoods with SpatialFDR < 0.01 and logFC| > 2 were considered significantly enriched or depleted. Significant neighborhoods were visualized on the embedding, and beeswarm plots were generated to show DA across clusters. For each cluster and comparison, condition-specific DA neighborhoods (COG52, COG53, WT) were reported and visualized using boxplots.

##### Functional enrichment analysis

Functional analysis of differentially expressed genes (DEGs) was conducted using QIAGEN Ingenuity Pathway Analysis (IPA) (1). A dataset comprising gene identifiers and corresponding expression values from apical progenitors, basal progenitors, and neurons, derived from organoids of the two mutated COG5 and a wild-type (WT) control, was uploaded into the IPA platform. Each gene identifier was mapped to its corresponding entity within the QIAGEN Knowledge Base. To identify significantly perturbed molecules, an absolute log fold change (logFC) threshold of ≥1 was applied. These molecules were then overlaid onto a global molecular network curated from the QIAGEN Knowledge Base. Biological function analysis was performed using a right-tailed Fisher’s Exact Test to calculate p-values, estimating the probability that the association between the DEGs and specific biological functions or diseases occurred by chance. A z-score was computed to predict the directionality (activation or inhibition) of each function or disease. Canonical pathway analysis was also carried out to determine the most significantly enriched pathways from the IPA library. Again, a right-tailed Fisher’s Exact Test was used to assess statistical significance, and a z-score was calculated to infer the likelihood of pathway activation or inhibition.

##### Trajectories Analysis

Trajectory inference was performed using Palantir (v1.4.4) separately for each condition of interest on Scanpy-preprocessed data. The top 2,000 highly variable genes were selected (Seurat v3), and diffusion maps were computed on Harmony-corrected PCA embeddings. As starting population, the apical progenitors (AP) cluster was defined separately for each condition, from which an initial cell was selected, while terminal states were identified automatically. Palantir was then run with 500 waypoints to infer pseudotime, differentiation potential (entropy), and terminal state probabilities, which were visualized on UMAP embeddings.

#### Organoids’ Glycolipidomics

Organoids at d40 were dissociated using the MACS tissue dissociation kit as described before and stored at -80°C for subsequent lipid glycomics analysis. Glycosphingolipid (GSL) analysis was performed according to the protocol for ganglioside analysis by LC-MS described by Jáñez Pedrayes et al. (2025). Briefly, cell pellets were thawed and resuspended in 250 µL of 1-butanol/methanol (1:1, v/v) with 1µM deuterated d5-GM3 (Avanti) as internal standard. Samples were sonicated and centrifuged for 15 min at 3,500 g. Pellets were processed for BCA assay. The supernatant was then purified using 0.20µm polytetrafluoroethylene (PTFE) filters (Millipore) and additional 500µL of 1-butanol/methanol (1:1, v/v) were filtered to ensure recovery of the GSLs. GSL solutions were evaporated in a SpeedVac and resuspended in 100µL of the chromatography starting conditions (methanol/water/isopropanol (70:15:15)) and transferred to MS vials. LC-MS analysis was performed on a Vanquish LC System (Thermo Scientific) equipped with a C18 column (Acquity UPLC-BEH, 130Å, 1. 7 μm; 2.1 mm x 150mm, Waters) coupled to an Orbitrap Fusion Lumos Tribrid mass spectrometer (Thermo Scientific) operating in negative ion mode (m/z range: 690-2000). The detailed MS protocol can be found in Jáñez Pedrayes et al. (2025). Data collection was performed using the Xcalibur software (Thermo Scientific). GSLs were identified and quantified according to their elution time and mass-to-charge (m/z) ratios using the analysis platform El-Maven for processing the data. GSL intensities were normalized to the internal standard and protein content for each sample.

#### Western Blot on human brain organoids

Organoids were collected and snap frozen in dry ice and stored at -80°C until use. Samples for western blot were prepared by adding SDS protein lysis buffer (2% SDS, 50mM TrisHCl pH8, 10mmM EDTA, 10% Glycerol) to the organoids (10% (w/v) lysate). Samples were boiled at 95°C for 2 minutes. To break all cell clusters the sample was passed through an insulin syringe, followed by ultrasonication (ultrasound time 3s, 10s interval, repeated from 5 to 15 times, ultrasonic power 40kW). Samples were then centrifuged for 10 minutes at 8000xg and the cell pellet was discarded. BCA assay kit (ThermoFisher, 23227) was used to determine protein concentration and the lysate was finally aliquoted and stored at -80°C until use.

Lysates corresponding to 20μg of protein were added with1X NuPAGE LDS Sample buffer (ThermoFisher, NP0007) and 1X NuPAGE^™^ Reducing Agent (ThermoFisher, NP0009). Samples were put at 70°C for 10 minutes to achieve heat-mediated denaturation.

Depending on target protein molecular weight, samples were loaded on the Nupage™ 3-8% Tris-Acetate Protein Gel (ThermoFisher, EA03785BOX) in 1X NuPAGE Tris Acetate Running Buffer (Thermofisher, LA0041) or Bolt™ Bis-Tris Plus Mini Protein Gels, 4-12% (ThermoFisher, NW04125BOX) in NuPAGE MES SDS Running Buffer (ThermoFisher, NP0002) together with the precision Protein Dual Color Standards (Bio-Rad, #1610374) ladder. For denaturing electrophoresis, the gel ran for 1 hour at 160 V constant. After the electrophoresis, proteins were transferred onto the 0.2 µm nitrocellulose membrane using Trans-Blot Turbo Mini Nitrocellulose Transfer Packs (Bio-Rad, 1704158) on a Trans-Blot Turbo TM Transfer System (Bio-Rad, 1704150) with parameters set to: 1.3A-25V for 10 minutes. The nitrocellulose membrane was washed with TBST (TBST: 10mM Tris-HCl, pH 8.0, 150 mM NaCl, 0.05% Tween 20) once for 5 min at room temperature. To prevent non-specific binding, the membrane was blocked with freshly prepared 5% non-fat dried milk (Millipore, 70166) in TTBS for 1 hour on a shaking platform at room temperature. The membrane was then incubated with the primary antibody prepared in 5% milk with gentle agitation at 4°C overnight. The next day the membrane was washed three times for 5 min each with TBST and then incubated with secondary HRP antibody at 1:5000 in TBST-5% milk for 1h at RT. The membrane was washed three times again for 5 minutes each with TBST. The Clarity Max Western ECL Substrate (Bio-Rad, 1705062) or the Clarity Western ECL Substrate (Bio-Rad, 1705060) was added to the membrane for 1 minute and finally the membrane was imaged with a ChemiDoc MP imaging system (Bio-Rad). Band intensities were quantified using Image Lab software (Bio-Rad) and analysed by GraphPad Prism. All original entire blots are available under request.

#### Preparation of membrane and cytosol fractions from human brain organoids

Organoids were weighted and Tris 50mM buffer containing the Protease inhibitor cocktail was added at a ratio 10% (w/v). The organoids were homogenized on ice using a potter. The homogenate was spinned for 10 minutes at 1000xg at 4°C to pellet the nuclei and tissue debris. The post-nuclear supernatant (PNS) was recovered, placed in a fresh eppendorf tube and centrifuged for 30 minutes at 12000xg at 4°C to separate the cytosol from the membrane fraction. The supernatant containing the cytosol was collected in a fresh eppendorf tube whereas the pellet containing membrane was resuspended in 1mL of 50mM Tris pH 7.4 supplemented with protease inhibitor cocktail. Protein concentration was determined with BCA protein assay kit. Samples were stored at -80°C and subsequently processed for western blotting.

#### Statistics and reproducibility

Statistical analysis was performed using Prism (GraphPad software v10.6.1). To test for statistical significance, we used one-way or two-way ANOVA for multiple comparisons with Tukey post test. For each graph, the number of samples, statistical test and the p value are reported in the corresponding figure legends. Threshold of statistical significance was set at ≤ 0.05 (****p < 0.0001, ***p<0.001; **p<0.01; *p < 0.05). Data derived from different technical replicates (n individual organoids or cells or images) of independent biological replicates (differentiation experiments) are specified in each legend and are represented as violin plots with median and quartiles or as bar plots with mean and SEM or SD. For single-cell RNA sequencing we sequenced N=3 independent batches (differentiation) per condition (control, COG52, COG53); n= 8 pooled organoids/each batch at day 40.

## RESULTS

### COG5 deficiency disrupts Golgi organization and retrograde trafficking

Apical radial glial cells (aRGs) are highly polarized neural progenitors characterized by a distinctive intracellular organization, including a non-pericentrosomal Golgi distributed along the apical process, between the apical plasma membrane and the nucleus (Fig. 1A). Given the central role of the Golgi apparatus in intracellular trafficking and glycosylation and considering the involvement of the Conserved Oligomeric Golgi (COG) complex in vesicle tethering and Golgi organization (Fig. 1B), we investigated whether COG5 deficiency affects Golgi structure and function in neural progenitors.

**Figure 1.**
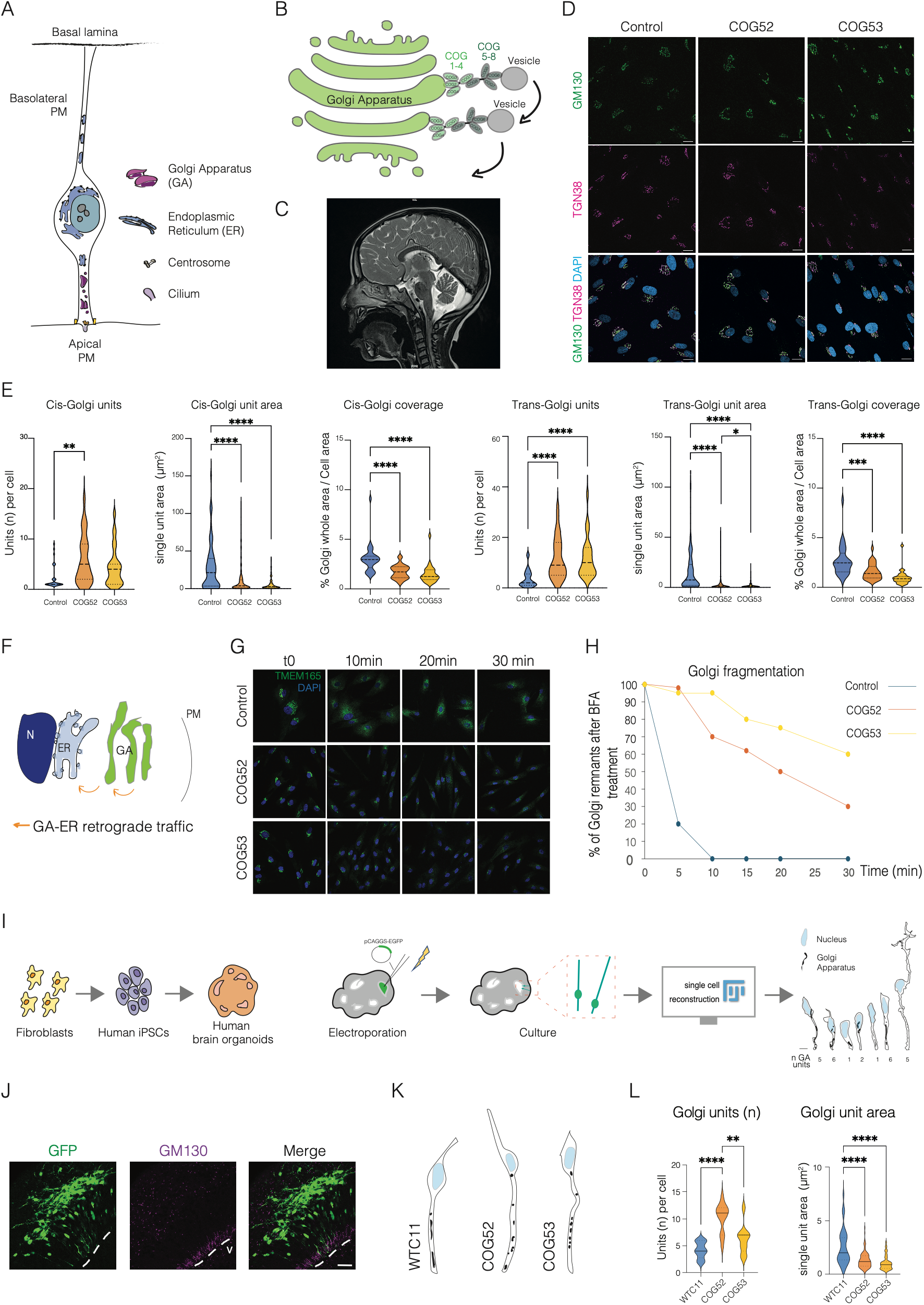
COG5 deficiency disrupts Golgi organization and retrograde trafficking. **(A)** Schematic representation of apical radial glial cell (aRG) morphology and intracellular organization, highlighting the polarized distribution of Golgi elements along the apical process. Abbreviations: GA=Golgi Apparatus, ER=Endoplasmic Reticulum, PM=Plasma Membrane. **(B)** Schematic representation of the Conserved Oligomeric Golgi (COG) complex and its role in vesicle tethering and retrograde trafficking within the Golgi apparatus. **(C)** MRI image from male patient (COG53) performed at age 4. **(D)** Representative immunofluorescence images of control and patient-derived fibroblasts stained for the cis-Golgi marker GM130, the trans-Golgi marker TGN38 and nuclei (DAPI). Scale bar 20 μm. **(E)** Quantification of number of cis- and trans-Golgi units and area of individual Golgi units, together with cis- and trans-Golgi whole area (sum of single units) normalized to cell area, in control and COG5-deficient fibroblasts. N=3 independent biological replicates. Total number of quantified cells per condition = 26-32. One-way ANOVA, Tukey post test, *p<0,05, **p<0,01, ***p<0,001 and ****p<0,0001. **(F)** Schematic representation of the Golgi-to-endoplasmic reticulum (GA–ER) retrograde trafficking pathway assessed by brefeldin A (BFA) treatment. Abbreviations: GA=Golgi Apparatus, ER=Endoplasmic Reticulum, PM=Plasma Membrane, N=Nucleus. **(G)** Representative immunofluorescence images of control (from unaffected mother) and COG5-deficient fibroblasts stained for Golgi marker TMEM165 and DAPI following BFA treatment at the indicated time points. **(H)** Quantification of Golgi fragmentation dynamics following BFA treatment, measured as percentage of Golgi remnants after BFA treatment. **(I)** Experimental workflow for the generation of patient-derived human brain organoids (see Material and Methods), sparse GFP labelling by electroporation, and single-cell reconstruction of apical radial glial cells. **(J)** Representative immunofluorescence image stained for GFP-labeled apical radial glial cell of control brain organoids to define cell outline and Golgi apparatus marker GM130 at day 40. Scale bar 20 μm. Dashed lines indicate ventricle surface and letter v. stans for ventricle. **(K)** Single-cell reconstructions of control and COG5-deficient apical radial glial cells showing Golgi distribution along the apical process. **(L)** Quantification of Golgi units number and Golgi single units area in control and COG5-deficient apical radial glial cells. N= 3 independent organoid batches and n=3 total apical progenitor cells analyzed across conditions for units; and N=2 n=41 for area. One-way ANOVA, Tukey post test, **p<0,01 and ****p<0,0001.

To first assess the relevance of the COG complex in neural development, we analyzed a publicly available mouse and human developmental transcriptomic dataset (Florio et al, 2015) and found that multiple COG subunits are expressed across neural progenitor and neuronal populations, including apical radial glia (aRGs), basal radial glia (bRGs), intermediate progenitors (IPCs), and neurons (Supplementary Fig. 1A-B). In both mouse and human datasets, COG subunits displayed broad expression across lineage progression, supporting a potential role for COG-dependent trafficking during neurogenesis.

We identified two patients of a novel COG5 mutation causative of Congenital Disorders of Glycosylation (CDG) and analyzed their derived fibroblasts (see Materials and Methods). The sibling patients, a female (COG52) and a male (COG53), were identified as carrying a novel homozygous intronic *COG5* variant associated with neurodevelopmental defects (Figure 1C; Supplementary Fig. 2A). They present a severe, progressive syndromic osteocondensing disorder, including growth retardation with microcephaly, cleft palate, profound bilateral sensorineural hearing loss, cerebellar and optic atrophy, and intellectual disability. An MRI performed at age 4½ revealed that the boy had cerebellar hypoplasia, primarily affecting the vermis, and a short, thin corpus callosum; stenosis of the internal auditory canals with poor visualization of the auditory nerves; a moderately dysplastic appearance of the vestibules; and a moderately thin appearance of the optic nerves (Figure 1C). Whole-genome sequencing identified a homozygous intronic variant in COG5 (NM_006348.3:c.632-33909T>A; p.?) in both affected children and the terminated fetus (Supplementary Figure 2A-B). The variant was present in the heterozygous state in the parents and in the unaffected sibling, consistent with recessive inheritance. RT–PCR analysis performed on lymphocytes from the two affected children demonstrated aberrant splicing, characterized by partial retention of intron 6 (Supplementary Figure 2C). This creates a frameshift and the introduction of a premature termination codon (NM_006348.3: p.(Asp211Valfs*12)), consistent with *in silico* predictions and resulting in partial preservation of spliced transcripts (Supplementary Figure 2C-D).

To investigate whether COG5 deficiency affects Golgi organization, we stained control and patient-derived fibroblasts for the cis-Golgi marker GM130 and the trans-Golgi marker TGN38 (Fig. 1D). Quantitative analysis revealed a marked increase in the number of cis- and trans-Golgi units in both patient lines, accompanied by a reduction in their individual size, consistent with a fragmented Golgi architecture (Fig. 1E, Supplementary Fig. 3A). Overall cell size was unchanged across conditions, indicating that the phenotype specifically affects Golgi organization rather than global cellular morphology (Supplementary Fig. 3B). In addition, the spatial distance between cis- and trans-Golgi compartments was increased in patient-derived fibroblasts (Supplementary Fig. 3C), further supporting altered Golgi organization.

Because the COG complex is primarily involved in retrograde trafficking, we next examined intracellular transport dynamics in fibroblasts (Fig. 1F; Supplementary Fig. 3D). Plasma membrane-to-Golgi trafficking assessed by Cholera Toxin B internalization (Supplementary Fig. 3D) was largely preserved, as indicated by comparable colocalization dynamics between Cholera Toxin B and cis- or trans-Golgi markers across conditions (Supplementary Fig. 3E–F). In contrast, Brefeldin A (BFA) treatment, a reversible inhibitor of trafficking between the endoplasmic reticulum (ER) and the Golgi, revealed delayed Golgi disassembly in COG5-deficient fibroblasts compared to controls, as assessed by time-course staining of the Golgi marker TMEM165 (Fig. 1G-H). The BFA causes dramatic Golgi disassembly with absorption of its components into the ER through a retrograde (backward) transport (Colanzi et al. 2013). These results indicate delayed Golgi disassembly driven by impaired GA- to-ER retrograde trafficking in COG5-deficient fibroblasts.

We next investigated whether similar Golgi defects occur in neural progenitors using human brain organoids from patient-derived induced pluripotent stem cells (iPSCs) (Supplementary Fig. 4A). Human brain organoids generated following the Lancaster protocol (Lancaster and Knoblich 2014) contain apical and basal progenitors organized in structures that mature and grow over time (Supplementary Fig. 4B-D).

Western blot analysis on patients’ organoids showed a marked reduction in COG5 protein levels (Supplementary Fig. 5). Given the role of COG5 within the COG complex, we also assessed the stability of interacting subunits revealing unchanged levels of the lobe A component COG4 and reduced lobe B subunit COG7 in both patients (Supplementary Fig. 5), indicating selective destabilization of lobe B components.

We then employed sparse GFP labelling by electroporation to reconstruct individual aRGs and visualize COG5 localization and Golgi organization at single-cell resolution (Fig. 1I). Using this approach, we observed that COG5 localizes to punctate Golgi structures positive for GM130 (Supplementary Fig. 4E), supporting COG5 association with the polarized Golgi network of apical radial glia cells (aRGs). In control organoids, Golgi stacks were distributed along the apical process, between the apical membrane and the nucleus (Fig. 1J–K), with the characteristic pattern previously described for rodent aRGs. Patient-derived aRGs displayed a significant increase in the number of Golgi stacks together with a reduction in their individual size (Fig. 1L), demonstrating a critical role for COG5 in preserving Golgi organization within neural progenitors.

Together, our results demonstrate that COG5 deficiency disrupt Golgi architecture via impaired retrograde trafficking, revealing a conserved Golgi disassembly (fragmentation) phenotype in both fibroblasts and apical radial glial cells.

### COG5 deficiency selectively impairs glycosphingolipid maturation in human brain organoids and accumulates sialylated N-glycans

Despite the severe neurodevelopmental phenotype observed in the patients, standard analyses of plasma N- and O-linked protein glycosylation (glycans linked to asparagine and glycans attached to serine or threonine residues, respectively) revealed no major abnormalities in transferrin or apoC-III glycosylation profiles (Supplementary Fig. 6A-B), indicating that these patients do not display the classical systemic glycosylation defects typically associated with COG-related CDGs. To the best of our knowledge, this is the first case of COG5-CDG patients without detectable serum protein glycosylation defects.

Although standard plasma transferrin and apoC-III glycosylation profiles were largely preserved, N-glycomic profiling of patient-derived brain organoids revealed that patient-derived organoids showed a slight increase in abundance of highly sialylated N-glycan species, indicating a tissue-specific remodelling of Golgi-dependent glycosylation rather than a classical loss of terminal sialylation (Fig. 2A).

**Figure 2.**
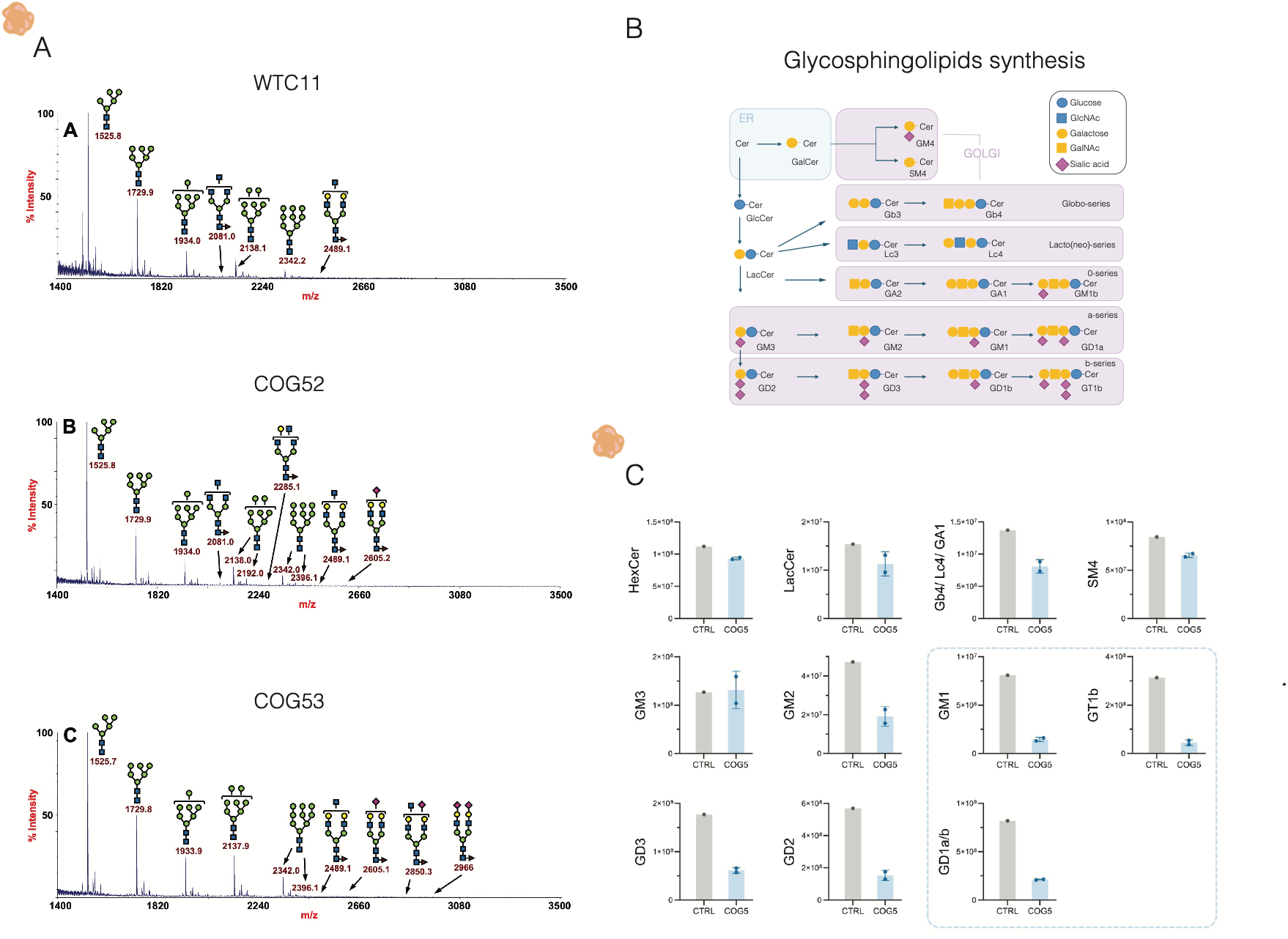
COG5 deficiency selectively enriches sialylated glycans and impairs glycosphingolipid (GSL) maturation in human brain organoids. **(A)** Representative MALDI-TOF mass spectra of N-glycans isolated from control, COG52, and COG53 brain organoids. Major N-glycan species are annotated according to their corresponding m/z values and predicted glycan structures. Comparison of the spectra reveals alterations in the relative abundance of complex N-glycan species in COG5-deficient organoids, including enrichment of highly sialylated glycans, indicating remodeling of Golgi-dependent N-glycan maturation despite the absence of major abnormalities in plasma glycosylation profiles. **(B)** Schematic representation of glycosphingolipid (GSL) biosynthesis within the endoplasmic reticulum (ER) and Golgi apparatus, illustrating the sequential maturation of ceramide-derived lipid species into complex gangliosides. **(C)** GSL relative abundances (MS intensities) in control (WTC11; n=1) and COG5-deficient organoids (n=2) dissociated at day 40. Each sample was generated by pooling 8 individual organoids together. Intensities are normalized to the internal standard and protein content per sample. Graphs show the average of the two predominant ceramide backbones, (d18:1/16:0) and (d18:1/18:0; d20:1/16:0). Graphs showing the intensities for each ceramide backbone separately can be found in supplementary figure 7. Most abundant ganglioside species in the brain are inside a dashed box. Data are represented as the mean ± SD. GD1a and GD1b are structural isomers with identical exact masses and are not chromatographically resolved, so they were measured as a combined signal by MS (referred to as GD1a/b). Gb4, Lc4 and GA1 have the same mass-to-charge (m/z) ratio and are measured together. GSL analysis reveals a reduction of mature, sialylated gangliosides in COG5-deficient organoids. HexCer: Hexosylceramide (Glucosylceramide + Galactosylceramide), LacCer: Lactosylceramide; SM4: sulfatide.

Given the central role of the Golgi apparatus in lipid glycosylation, we next investigated whether COG5 deficiency affects glycosphingolipid (GSL) maturation in neural tissue. We therefore performed lipidomic profiling on patient-derived brain organoids at day 40, focusing on GSL species, known to be involved in membrane organization and signalling (Fig. 2B).

COG5-deficient organoids displayed a marked reduction in complex, sialylated gangliosides (Fig. 2C). Species-level analysis further demonstrated that these alterations occur across multiple ceramide backbones, including d18:1/16:0 and d18:1/18:0 species, indicating a broad impairment of glycosphingolipid maturation rather than selective alterations of individual lipid classes (Supplementary Fig. 7).

This finding contrasts with the slight increase in the abundance of sialylated N-glycans, and suggests that COG5 deficiency differentially affects protein- and lipid-linked glycosylation pathways in neural tissue.

### COG5 deficiency alters aRG lineage progression and delays neurogenesis

Consistent with the essential role of Golgi apparatus in brain development, the majority of CDG patients present brain-related symptoms such as microcephaly, hypotonia, psychomotor retardation, delayed speech development and cerebellar atrophy (D’Souza et al. 2020). Given the prominent Golgi defects observed in aRGs and the patients’ neurodevelopmental defects, we next investigated whether COG5 deficiency affects progenitor dynamics and neurogenesis in the neurodevelopmental model of human brain organoids.

Morphometric analysis revealed that patient-derived organoids were reduced in size compared to controls, with a more pronounced decrease observed in the COG53 patient line, together with fewer ventricular-like structures in both patient-derived organoids, indicating altered tissue organization (Fig 3A-B). In contrast, individual ventricle area and apical surface length were largely unchanged across conditions (Fig. 3B). To determine whether macroscopic observations were paralleled by changes in tissue organization, we quantified ventricular architecture across developmental stages and observed that total apical-to-basal thickness was reduced in both COG5 organoids (Fig. 3C). This reduction was accompanied by a decreased thickness of the neuronal layer, particularly in COG53 patients’ organoids, whereas the ventricular zone was not affected.

**Figure 3.**
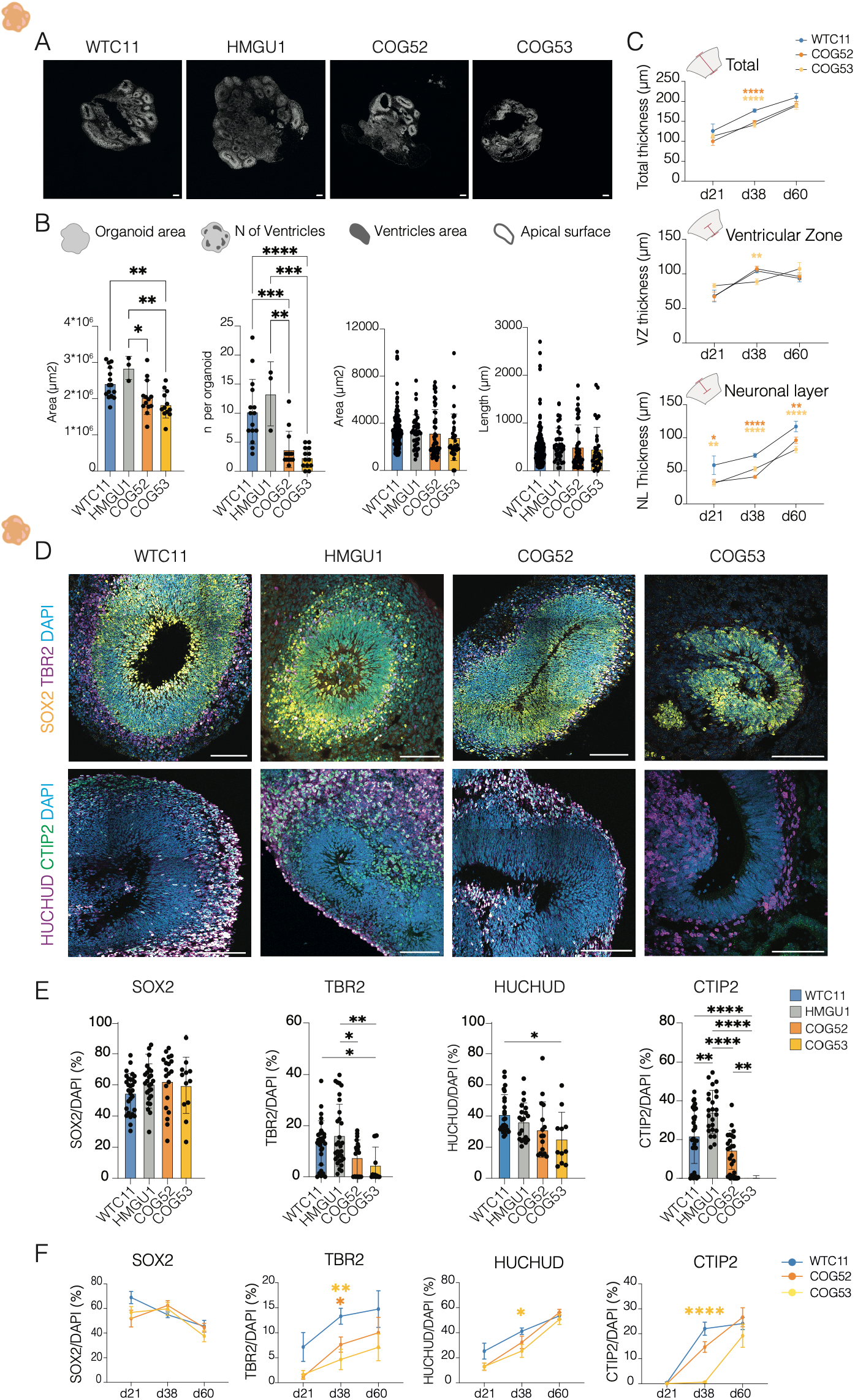
COG5 deficiency alters organoid architecture and neural progenitor progression. **(A)** Representative immunofluorescence images of control (WTC11, HMGU1) and patient-derived (COG52, COG53) human brain organoids stained for DAPI at day 38. Scale bar 200 μm. **(B)** Quantification of organoid area, and ventricular-like structures’ parameters: number, area, and apical surface length, in control and patient-derived organoids. N= 3 independent organoid batches and n=39 total organoids analyzed across conditions. One-way ANOVA, Tukey post test, p<0,05, **p<0,01, ***p<0,001 and ****p<0,0001. **(C)** Quantification of total ventricular thickness, ventricular zone (VZ) thickness, and neuronal layer (NL) thickness across developmental stages (days 21, 38, and 60) of control and COG5 organoids. N= 3 independent organoid batches and n=3 total organoids/batch per each timepoint and condition. Two-way ANOVA, Tukey post test, *p<0,05, **p<0,01, ****p<0,0001. **(D)** Representative immunofluorescence images of organoid stained for SOX2, TBR2, HuC/HuD, CTIP2, and DAPI, identifying apical progenitors, basal progenitors, and neuronal populations at day 38. Scale bar 20μm. **(E)** Quantification of percentage of SOX2⁺ apical progenitors, TBR2⁺ basal progenitors, HuC/HuD⁺ and CTIP2⁺ neurons in control and COG5-deficient organoids at day 38. N= 3 independent organoid batches and n=3 total organoids/batch per condition. One-way ANOVA, Tukey post test, *p<0,05, **p<0,01, and ****p<0,0001. **(F)** Bottom panels show temporal analysis of these populations across developmental stages (days 21, 38, and 60). N= 3 independent organoid batches and n=3 total organoids/batch per each timepoint and condition. Two-way ANOVA, Tukey post test, *p<0,05, **p<0,01.

To determine whether these structural alterations were associated with changes in neural populations, we analyzed organoids for markers of apical progenitors (SOX2), basal progenitors/IPC (TBR2), and neurons (HuC/HuD and CTIP2) (Fig. 3D). While the proportion of SOX2⁺ apical progenitors was largely preserved, TBR2⁺ basal progenitors/IPC were significantly reduced in both patient lines. Neuronal populations were also affected, with reduced HuC/HuD⁺ neurons and a marked decrease in CTIP2⁺ neurons, particularly in COG53 patient organoids (Fig. 3E).

We next assessed whether these changes reflected altered developmental timing by quantifying these progenitor and neuronal populations across different stages of neurogenesis (days 20, 40, and 60). Whereas SOX2⁺ apical progenitors and HuC/HuD⁺ neurons followed similar trajectories across conditions, TBR2⁺ progenitors and CTIP2⁺ neurons showed major defects at day 40, corresponding to the peak of the neurogenic stage of the organoid model, and accumulated more slowly in patient-derived organoids (Fig. 3F).

Together, these findings indicate that disruption of Golgi organization in aRGs alters lineage progression and delays the generation and maturation of downstream progenitor and neuronal populations.

### COG5 deficiency alters developmental trajectories and promotes premature neuronal differentiation

To investigate how COG5 deficiency affects the generation and maturation of neural cell types, we performed single-cell RNA sequencing on day 40 organoids. Clustering analysis identified major neural and non-neural populations, including apical progenitors (APs), basal progenitors (BPs), intermediate progenitor/neuron populations (IPC_N), neuronal clusters (Imm_N, N1, N2), glial progenitors (Glia_P), extracellular matrix-enriched progenitors (P_ECM), and choroid plexus cells (ChP) (Fig. 4A–B; Supplementary Fig. 8A–B).

**Figure 4.**
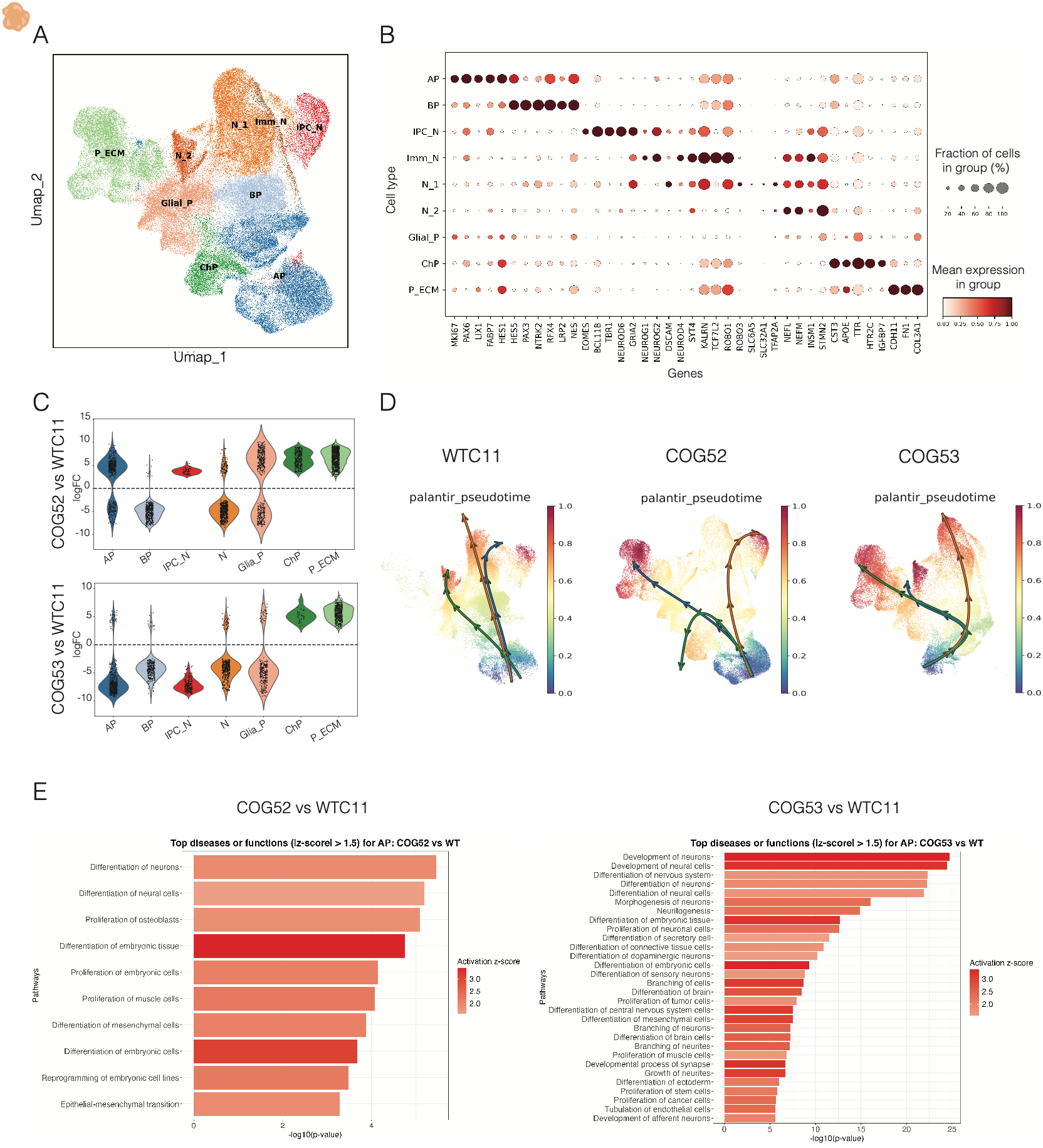
COG5 deficiency alters developmental trajectories and promotes premature neuronal differentiation. **(A)** UMAP representation of single-cell transcriptomic profiles from control and COG5-deficient organoids, identifying major neural and non-neural populations, including apical progenitors (APs), basal progenitors (BPs), intermediate progenitor/neuron populations (IPC_N), neuronal clusters (N_1, N_2, Imm_N), extracellular matrix-enriched progenitors (P_ECM), glial progenitors (Glial_P), and choroid plexus cells (ChP). **(B)** Dot plot showing expression of representative marker genes used for cluster annotation across the identified cell populations. Dot size represents the fraction of expressing cells, and colour intensity indicates relative mean expression levels. **(C)** Differential abundance analysis comparing COG52 and COG53 organoids with wild-type controls, showing depletion of basal progenitors and neuronal populations together with enrichment of choroid plexus and extracellular matrix-associated cell states. **(D)** Pseudotime reconstruction of developmental trajectories in Palantir, using apical progenitors as the starting population. Control organoids follow canonical neurogenic routes through intermediate progenitor states, whereas COG5-deficient organoids display altered trajectories characterized by direct differentiation toward neuronal states and diversion toward extracellular matrix-associated populations. **(E)** Top Diseases and Biological Functions identified by Ingenuity Pathway Analysis (IPA) of differentially expressed genes in apical progenitors of COG52 and COG53 organoids compared with controls. Embryonic and neuronal differentiation pathways are enriched in both patient lines, with a stronger activation signature in COG53 organoids.

Differential abundance analysis (DA) to evaluate cell type composition, revealed depletion of basal progenitors and neuronal populations in patient-derived organoids, together with enrichment of extracellular matrix-associated cell states and choroid plexus, (Fig. 4C; Supplementary Fig. 8C).

To further define lineage progression, we reconstructed pseudotime trajectories using APs as the starting population. While control organoids followed canonical neurogenic routes, patient-derived organoids displayed altered trajectories with a reduction of indirect neurogenesis and an increase of direct neurogenesis in COG patient organoids. In addition, COG patient organoids display an increase in a cell population associated with extracellular matrix production (Fig. 4D).

Consistent with these observations, transcriptional alterations were already present at the early stage of apical progenitors (Supplementary Fig. 8D). In line with previously described trafficking defects, shared DEGs between COG patient organoids vs controls, show downregulation of IRAG2, an endoplasmic reticulum–associated type II transmembrane protein involved in intracellular organization and trafficking. Moreover, genes involved in neuronal differentiation and connectivity such as CNTN4, SLITRK2, and TMEM132C were significantly upregulated. Conversely, genes associated with developmental regulation, including DLX6-AS1 and RIPPLY2 whose expression is regulated by Notch and Wnt pathways (Dunty et al. Development, 2008), were downregulated. Pathway analysis on apical progenitors further revealed activation of multiple differentiation programs, including neuronal pathways such as “differentiation of neurons” and “neuritogenesis” together with embryonal and mesenchymal tissues, with a stronger effect in COG53 patient organoids (Fig. 4E). These differential expression patterns support the transcriptional shift from progenitor-associated programs toward differentiation observed in COG5-deficient organoids. Together, these findings indicate that COG5 deficiency rewires developmental trajectories, impairing progenitor lineage progression and promoting premature differentiation.

### COG5 deficiency disrupts NOTCH1 localization and signalling

Our single-cell transcriptomic analyses identified widespread perturbations in pathways regulating neural progenitor maintenance and differentiation. Among these, the Notch pathway emerged as a prominent candidate, with both COG5-deficient organoid lines exhibiting reduced expression of canonical Notch target genes, including members of the HES family (Fig. 5A). These findings suggested impaired Notch signalling as a potential mechanism underlying the altered lineage progression observed in COG5-deficient organoids. Notably, a similar reduction in Notch activity has been associated with premature neuronal differentiation in human cortical models carrying NOTCH2NL loss-of-function mutations (Fiddes et al. 2018).

**Figure 5.**
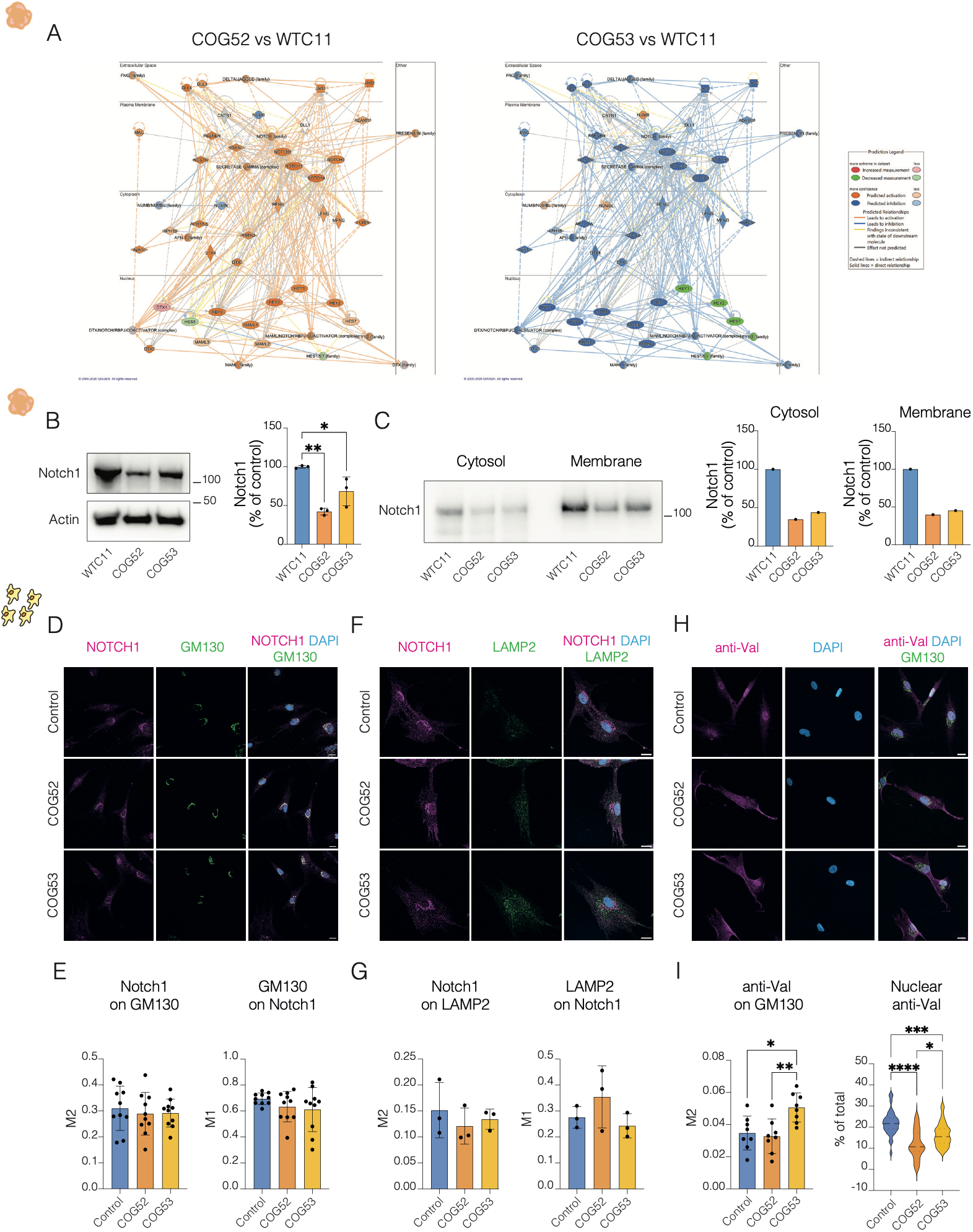
COG5 deficiency is associated with impaired Notch signalling and altered ACBD3 localization. **(A)** Ingenuity Pathway Analysis (IPA) of the NOTCH signalling network in apical progenitors of COG52 and COG53 organoids compared with wild-type controls, highlighting altered expression of Notch signalling targets in COG5-deficient organoids. **(B)** Western blot analysis and quantification of NOTCH1 expression levels in control and COG5-deficient organoids. N=3 independent organoid batches and n=3 total organoids/batch per each condition. One-way ANOVA, Tukey post test, *p<0,05, **p<0,01. **(C)** Western blot analysis and quantification of membrane and cytosol fractions of NOTCH1 expression levels in control and COG5-deficient organoids. N=1 independent organoid batches and n=3 total organoids per each condition. **(D-F-H)** Representative immunofluorescence images of total Notch1, GM130, LAMP2, and Cleaved Notch1 (Val1744), in control and COG5-deficient organoids. Scale bar 20μm. (**E-G-I**) Colocalization quantified as M1 or M2 value from JacoP plugin in Fiji and percentage of Val staining in the nucleus. N=1 independent experiment and n=3-10 total images quantified per each condition. One-way ANOVA, Tukey post test, *p<0,05, **p<0,01, ***p<0,001 and ****p<0,0001.

To determine whether these transcriptional changes were accompanied by altered receptor abundance, we examined NOTCH1 protein levels in brain organoids. Western blot analysis demonstrated a reduction in full-length NOTCH1 in both COG52 and COG53 organoids compared with controls (Fig. 5B). Subcellular fractionation further revealed that both membrane-associated and cytosolic NOTCH1 were markedly decreased (Fig. 5C).

Given the central role of Golgi trafficking in NOTCH1 maturation and activation, we next investigated the intracellular localization of the receptor in patient-derived fibroblasts. NOTCH1 remained associated with the Golgi apparatus, with no significant changes in colocalization with the cis-Golgi marker GM130 between conditions (Fig. 5D,E) and showed comparable localization with the lysosomal marker LAMP2 (Fig. 5F,G), arguing against enhanced lysosomal targeting as the primary cause of reduced receptor abundance.

We then examined the localization of the activated form of NOTCH1 using an antibody recognizing the cleaved Val1744 epitope. Although activated NOTCH1 remained associated with the Golgi, its colocalization with GM130 was significantly increased in COG53 fibroblasts (Fig. 5H,I; Supplementary Fig. 8A). In parallel, the nuclear pool of cleaved NOTCH1 (Notch Intra Cellular Domain, NICD) was markedly reduced in both COG5 deficient fibroblasts, consistent with impaired accumulation of the signalling-competent receptor in the nucleus (Fig. 5I). Together, these findings indicate that COG5 deficiency perturbs the intracellular distribution of activated NOTCH1, providing a mechanistic explanation for the reduced Notch transcriptional and molecular output observed in patient-derived organoids.

### Golgi disorganization redistributes ACBD3 and is associated with altered progenitor division

To investigate how disruption of Golgi organization might affect Notch signalling, we focused on ACBD3, a Golgi-associated scaffold protein that regulates asymmetric cell division by controlling the interaction between NUMB and NOTCH during neural progenitor mitosis (Zhou et al. 2007).

Immunoblot analysis demonstrated comparable ACBD3 protein levels in control and COG5-deficient organoids (Fig. 6A), indicating that COG5 deficiency does not alter ACBD3 abundance. In contrast, immunofluorescence analyses revealed a significant reduction in ACBD3 colocalization with the cis-Golgi marker GM130 (Fig. 6B,C), indicating spatial redistribution away from the Golgi apparatus.

**Figure 6.**
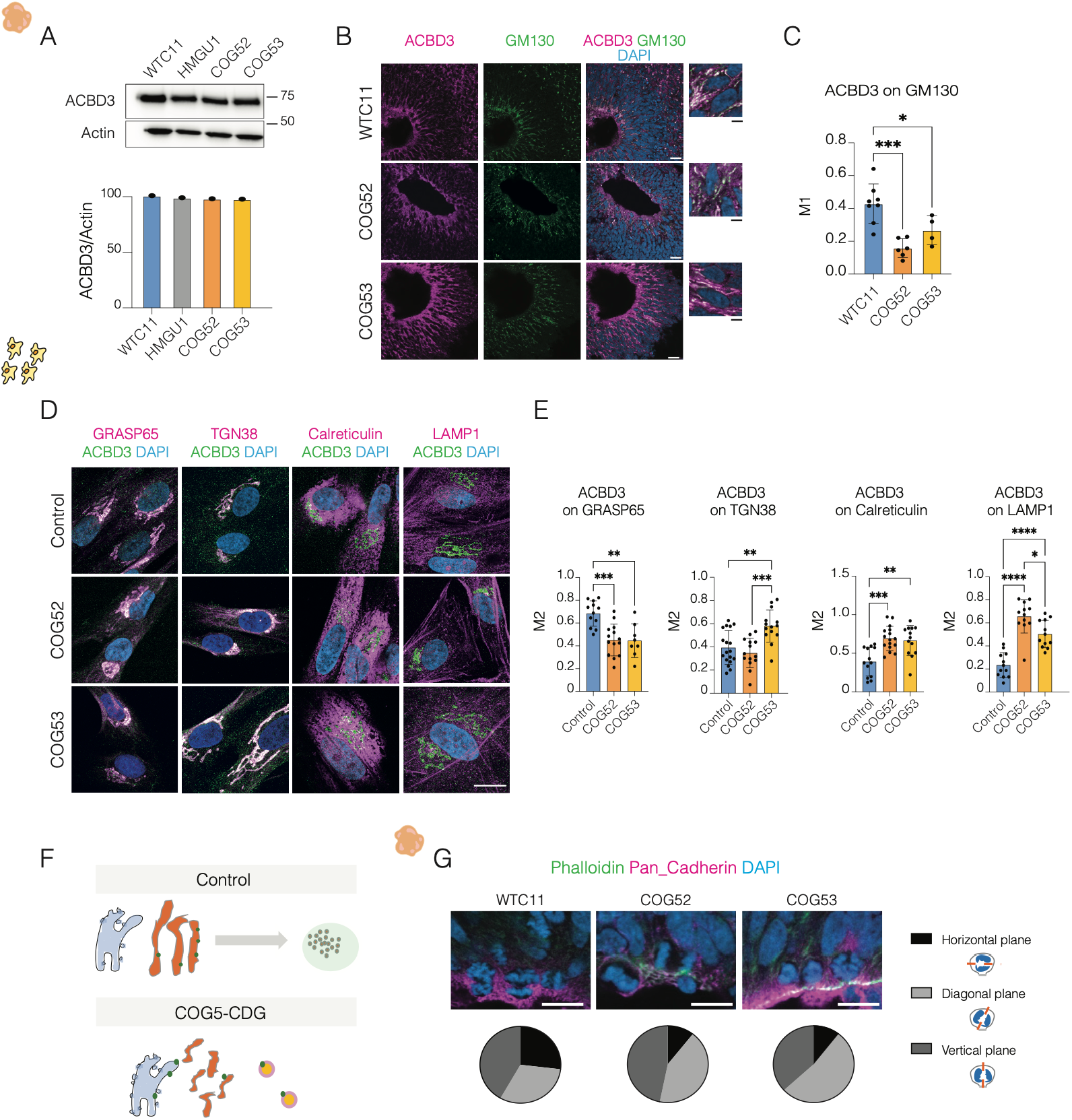
Altered ACBD3 localization and cleavage plane orientation in COG5-deficient samples. **(A)** Western blot analysis and quantification of total ACBD3 protein levels in control and COG5-deficient organoids. N=1 independent organoid batches and n=3 total organoids/batch per each condition. **(B)** Representative immunofluorescence images of ACBD3 and the Golgi marker GM130 in control and patient-derived organoids at day40. Scale bar 20 and 4 μm. **(C)** Quantification of ACBD3 colocalization with GM130, as M1 value from JacoP plugin in Fiji. N= 4-8 total images quantified per each condition, from 3 independent organoid batches (3 pooled organoids/batch). One-way ANOVA, Tukey post test, *p<0,05, **p<0,01. **(D)** Representative images of ACBD3 localization relative to intracellular compartments, including cis-Golgi (GRASP65), trans-Golgi (TGN38), endoplasmic reticulum (Calreticulin), and lysosomes (LAMP1) in control and COG5_deficient fibroblasts. Scale bar 20μm. **(E)** Colocalization quantified as M1 value from JacoP plugin in Fiji. N=2 independent experiments and n=12-18 total images quantified per each condition. One-way ANOVA, Tukey post test, *p<0,05, **p<0,01, ***p<0,001 and ****p<0,0001. **(F)** Working model illustrating the proposed redistribution of ACBD3 from Golgi compartments to alternative intracellular membrane compartments in COG5-deficient cells. **(G)** Analysis of cleavage plane orientation in mitotic apical radial glial cells from brain organoids stained for Phalloidin and Pan-cadherin. Representative images and quantification of horizontal (angle 0°-30°), diagonal (angle 30°-60°), and vertical (angle 60°-90°) divisions are shown for control and COG5-deficient organoids. Scale bar 20μm. Pie chart quantification for percentage of divisions type. N=3 independent organoid batches and n= total mitotic divisions quantified for WTC11=67, COG52=47, COG53=38. Exact values as follows: WTC11 H:27%, D:31%, V:41%; COG52 H:10%, D:43%, V:47%; COG53 H:11%, D:53%, V:37%

To define its altered intracellular localization, we examined ACBD3 colocalization with markers of distinct membrane compartments. In both COG5-deficient fibroblasts, ACBD3 showed reduced association with the Golgi marker GRASP65 and increased colocalization with the endoplasmic reticulum marker Calreticulin and the lysosomal marker LAMP1, whereas its association with the early and late endosomal markers RAB5 and RAB7 remained largely unchanged (Fig. 6D,E; Supplementary Fig. 9B,C). These findings indicate that loss of COG5 alters the spatial organization of ACBD3 rather than its expression, consistent with a broader disruption of Golgi-dependent membrane organization (Fig. 6F).

Because ACBD3 contributes to the regulation of asymmetric neural progenitor divisions through the Notch pathway, we next examined mitotic cleavage orientation in apical radial glial cells. Compared with controls, both COG5-deficient organoids exhibited fewer horizontal divisions and an increase in diagonal cleavage planes (Fig. 6G), consistent with altered daughter-cell fate decisions and the premature depletion of the progenitor pool.

Together, these findings support a model in which COG5 deficiency leads to disruption of Golgi organization, redistribution of the ACBD3 signalling platform, altered intracellular handling of activated NOTCH1, and impaired Notch signalling, thereby linking organelle architecture to neural progenitor fate during human corticogenesis.

## DISCUSSION

Our study identifies Golgi organization as a critical regulator of apical radial glial cell (aRG) behaviour and provides a biological framework linking COG5 deficiency to altered human brain development. By combining patient genetics, human brain organoids, glycolipidomics, and single-cell transcriptomics, we show that disruption of COG5-dependent Golgi organization affects lineage progression through coordinated defects in intracellular trafficking, membrane glycosylation, and the spatial organization of developmental signalling. A central finding of our study is that COG5 deficiency disrupts the distinctive Golgi organization of aRGs. Indeed, unlike most cell types, aRGs display a highly polarized intracellular architecture in which Golgi elements are distributed along the apical process (Taverna et al. 2016). We show that COG5 loss leads to fragmentation of these Golgi compartments in both fibroblasts and neural progenitors, indicating that COG5 is required for maintaining Golgi organization across different cellular contexts. Importantly, our study in patient fibroblasts revealed that these structural alterations result from impaired Golgi-to-ER retrograde trafficking, consistent with the established role of the COG complex in vesicle tethering, delivery of vesicle’s content and membrane recycling (Blackburn et al. 2019).

At the developmental level, disruption of Golgi organization is associated with altered lineage progression in human brain organoids. We observe reduced basal progenitor abundance, increased direct neurogenesis, and decreased indirect neurogenesis ultimately leading to a decreased production of neurons, indicating that Golgi dysfunction impacts key developmental transitions downstream of aRGs. Single-cell transcriptomic analysis further reveals rewired developmental trajectories characterized by reduced expression of progenitor-associated transcriptional programs and premature differentiation from apical progenitors. Together, these observations suggest that proper Golgi organization is required to coordinate the balance between progenitor maintenance and neuronal differentiation.

A particularly intriguing finding of our study is the differential impact of COG5 deficiency on protein- and lipid-linked glycosylation. While classical COG-related CDGs are typically diagnosed through abnormalities in serum glycosylation assays (Mól et al. 2026), the patients described here showed no major alterations in transferrin or apoC-III glycosylation. In contrast, glycomic and lipidomic analyses of patient-derived brain organoids revealed selective remodelling of Golgi-dependent glycosylation, characterized by a slight increase in the abundance of highly sialylated complex N-glycans together with a marked reduction in mature gangliosides. These findings indicate that COG5 deficiency does not uniformly impair Golgi glycosylation, but differentially affects protein- and lipid-linked glycosylation in a tissue-specific manner.

One possible explanation for the uncoupling between protein and lipid glycosylation is that glycoproteins and glycolipids traverse partially distinct trafficking and maturation routes within the Golgi apparatus. Rather than operating as a single homogeneous processing unit, the Golgi is increasingly recognized as a modular network composed of functionally specialized compartments, or distinct mini-stacks, that expose different cargo classes to tailored enzymatic micro-environments (Harada et al. 2024). Under this paradigm, N-glycosylation and glycolipid biosynthesis may rely on spatially segregated sub-domains whose specific enzyme and acceptor pools are continuously defined by COG-dependent retrograde trafficking. Furthermore, specialized recycling pathways, such as the retrieval of plasma membrane glycoconjugates for terminal sialylation in distinct Golgi sub-compartments, could render specific glycosylation pathways disproportionately vulnerable to COG dysfunction. This spatial and functional segregation may be particularly critical in highly polarized cells like neural progenitors, where precise local control of membrane composition governs developmental signalling. Interestingly, the predominance of high-mannose N-glycans in wild-type human brain organoids is consistent with recent evidence that neurons rely extensively on unconventional secretory pathways (Hanus et al. 2016). Under these conditions, glycoproteins and glycolipids may exhibit distinct sensitivities to perturbations of Golgi organization, potentially explaining why membrane glycolipid homeostasis appears particularly vulnerable to COG5 deficiency.

The selective remodelling of glycosphingolipids observed in COG5-deficient organoids is likely to have consequences beyond membrane composition. Gangliosides are major constituents of membrane microdomains that regulate receptor organization, trafficking, and signalling (Hirabayashi 2012). Alterations in ganglioside homeostasis may therefore contribute to the broader signalling defects identified by our transcriptomic analyses. Consistent with this possibility, among identified developmental pathways, Notch signalling showed reduced expression of canonical downstream targets including members of the HES family.

At the protein level, we further demonstrate reduced NOTCH1 abundance together with altered intracellular distribution of the receptor. While NOTCH1 remained associated with the Golgi apparatus and did not exhibit increased lysosomal localization, the activated receptor displayed increased Golgi association and reduced nuclear accumulation, consistent with altered signalling output. In parallel, the Golgi-associated scaffold protein ACBD3 was redistributed away from Golgi compartments toward endoplasmic reticulum and lysosomal membranes without changes in overall protein abundance. Given the established role of ACBD3 in coordinating NUMB-dependent regulation of Notch signalling during asymmetric progenitor division (Zhou et al., 2007), these findings suggest that disruption of Golgi organization alters the spatial organization of the Notch signalling machinery rather than simply affecting receptor abundance. Although the functional significance of ACBD3 redistribution remains to be fully established, its re-localization may reflect a broader perturbation of endomembrane organization associated with COG deficiency (Willett et al. 2014).

Consistent with altered Notch pathway activity, COG5-deficient organoids exhibited changes in cleavage plane orientation, with fewer horizontal and more diagonal divisions of apical radial glial cells. Because cleavage orientation influences the balance between symmetric progenitor expansion and asymmetric neurogenic divisions, these defects provide a plausible cellular mechanism linking altered Golgi organization to the premature lineage progression observed in COG5-deficient organoids. Importantly, however, our findings do not establish a direct causal relationship between Golgi dysfunction and Notch signalling. Rather, they support a model in which Notch represents one component of a broader signalling network affected by disruption of Golgi architecture, altered membrane trafficking, and remodelling of membrane glycosylation. Future studies will be required to determine how these Golgi-dependent processes cooperate to coordinate developmental signalling during human corticogenesis.

In conclusion, our study establishes a multi-level connection between Golgi organization, intracellular trafficking, membrane glycosylation, and developmental signalling in human neural progenitors. Our findings support a broad model in which Golgi architecture functions as an organizer of signalling environments that govern progenitor behaviour and cell fate decisions during human corticogenesis. Collectively, this study identifies intracellular organization as an additional regulatory layer of cortical development and suggests that disruption of organelle architecture can directly influence developmental trajectories in the developing human brain.

## FIGURES LEGENDS

**Supplementary Figure 1.**
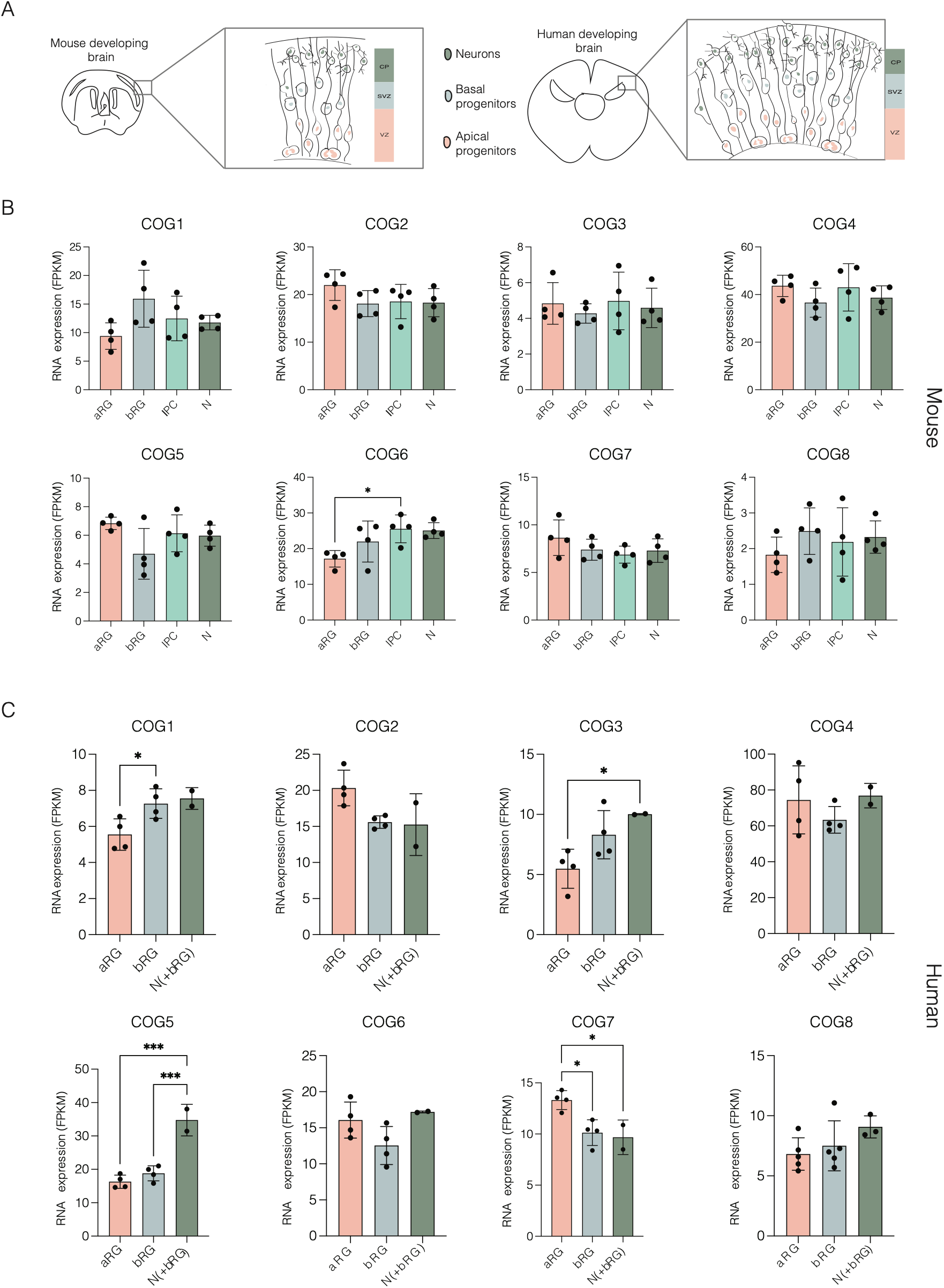
Expression of COG complex subunits across neural progenitor and neuronal populations. **(A)** Schematic representation of the developing mouse and human cortex highlighting major neural cell populations at mid neurogenesis, including neuroepithelial cells (NEC), apical radial glia (aRG), basal radial glia (bRG), intermediate progenitors (IPC), and neurons (N). **(B)** RNA expression levels of COG complex subunits (COG1–COG8) across neural cell populations in the developing mouse cortex at E14.5. **(C)** RNA expression levels of COG complex subunits (COG1–COG8) across neural populations in the developing human cortex at 13 postconceptional week (pcw). Data from GSE65000 mouse and human fpkm_matrix (Florio et al., 2015). N=2-4 independent replicates per cell populations. One-way ANOVA, Tukey post test, *p<0,05, **p<0,01, ***p<0,001.

**Supplementary Figure 2.**
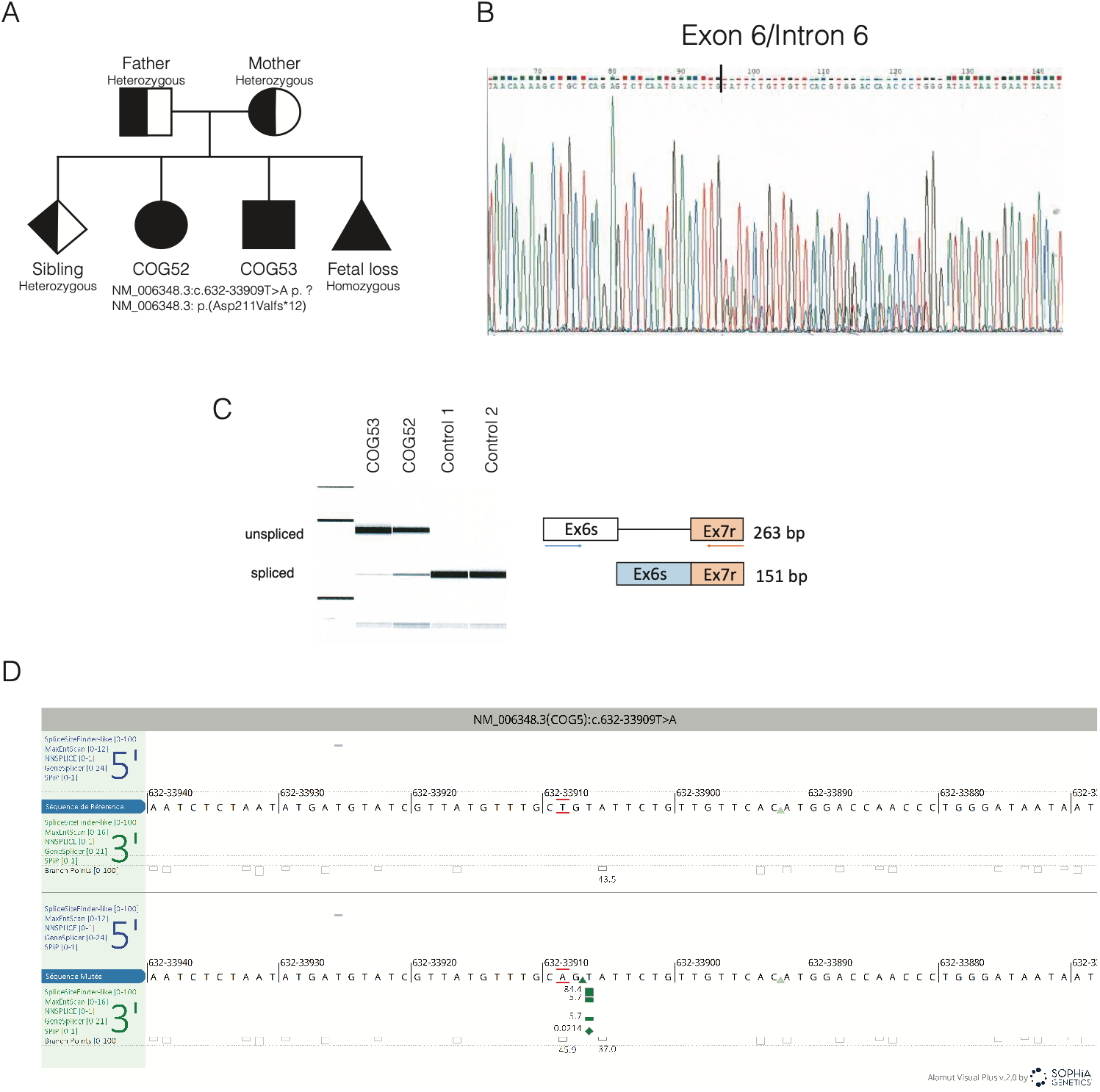
Genetic characterization of COG5-CDG patients. **(A)** Pedigree diagram of affected individuals and details of homozygous intronic COG5 variant (NM_006348.3:c.632-33909T>A). The family presents two affected children (male and female), one affected fetus, and one unaffected heterozygous child. **(B)** Sanger sequencing validation of the splice-altering mutation and resulting frameshift predicted to generate the truncated protein p.(Asp211Valfs*12). **(C)** Electrophoresis gel of COG5 RNA product showing that the mutation causes aberrant splicing with retention of intron 6 of COG5 gene in COG52 and COG53 patients. Schematic representation of the mutation, resulting in partial retention of intron 6 between exon 6 and exon 7 and generation of a frameshifted transcript **(D)** Details of homozygous intronic COG5 variant (NM_006348.3:c.632-33909T>A).

**Supplementary Figure 3.**
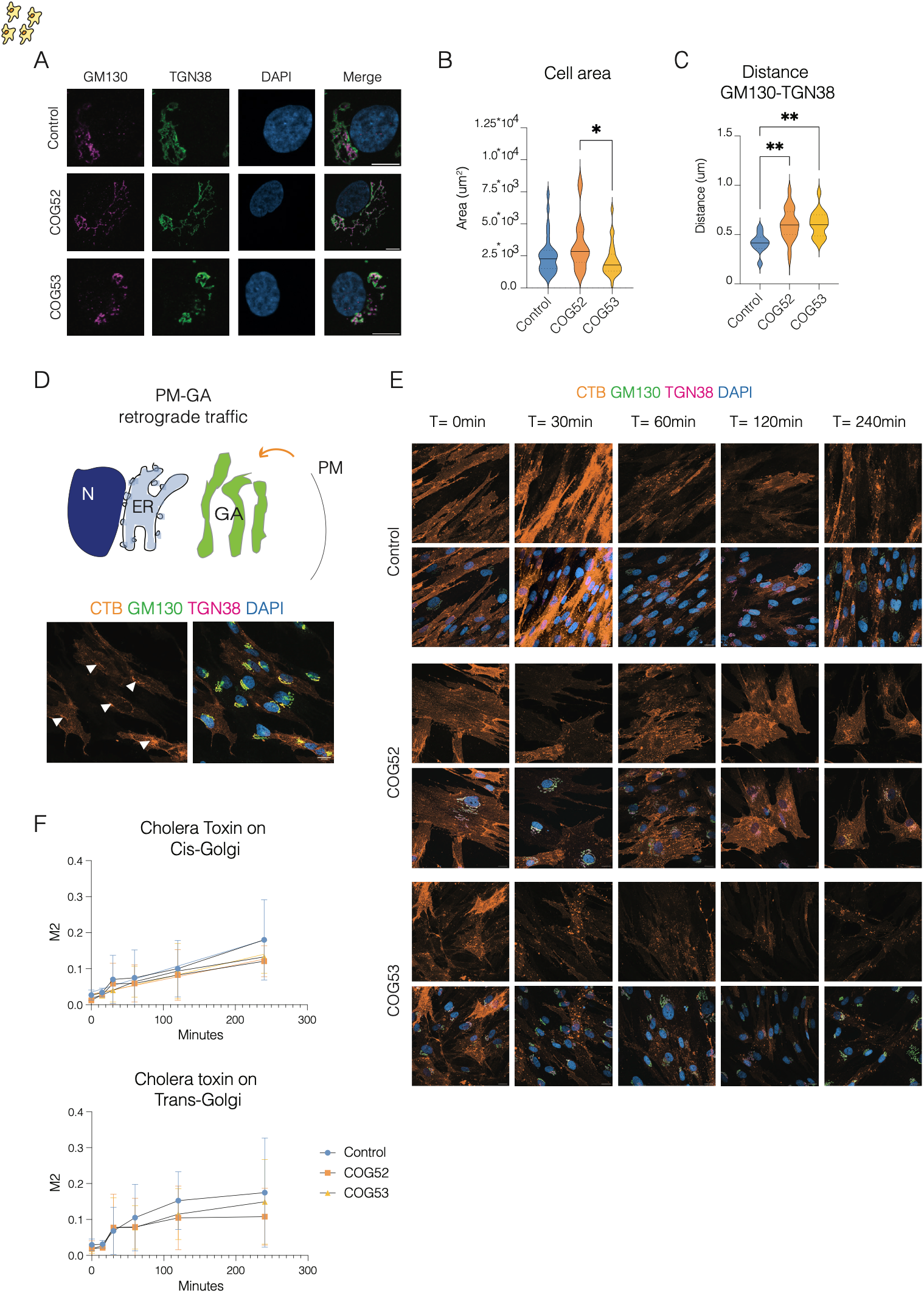
Golgi organization and plasma membrane-to-Golgi trafficking in COG5-deficient fibroblasts. **(A)** Representative immunofluorescence images of control and COG5-deficient fibroblasts stained for the cis-Golgi marker GM130, the trans-Golgi marker TGN38, and DAPI. Scale bar 20 μm. **(B)** Quantification of cell area in control and patient-derived fibroblasts. N=3 independent biological replicates. Total number of quantified cells: WT=32, COG52=29, COG53=27. One-way ANOVA, Tukey post test. **(C)** Quantification of the spatial distance between cis- and trans-Golgi compartments in control and patient-derived fibroblasts. N=1 independent biological replicate. Total number of quantified cells: WT=10, COG52=40, COG53=12. One-way ANOVA, Tukey post test, **p<0,01. **(D)** Schematic representation of intracellular retrograde trafficking route from plasma membrane-to Golgi and representative image of colocalization (highlighted by white arrows) between Cholera Toxin B (CTB) and Golgi markers. **(E)** Representative images from Cholera Toxin B (CTB) pulse–chase experiment showing CTB internalization and trafficking to Golgi compartments over time in control and COG5-deficient fibroblasts. Staining for CTB, GM130, TGN38, and DAPI. Scale bar 20 μm **(F)** Quantification of CTB colocalization with cis-Golgi (GM130) and trans-Golgi (TGN38) markers, performed by JaCoP plugin in Fiji, in control and COG5-deficient fibroblasts. N=3 independent biological replicates. n = 3-4 images per timepoint. Data are shown as mean ± SEM. One-way ANOVA, Tukey post test.

**Supplementary Figure 4.**
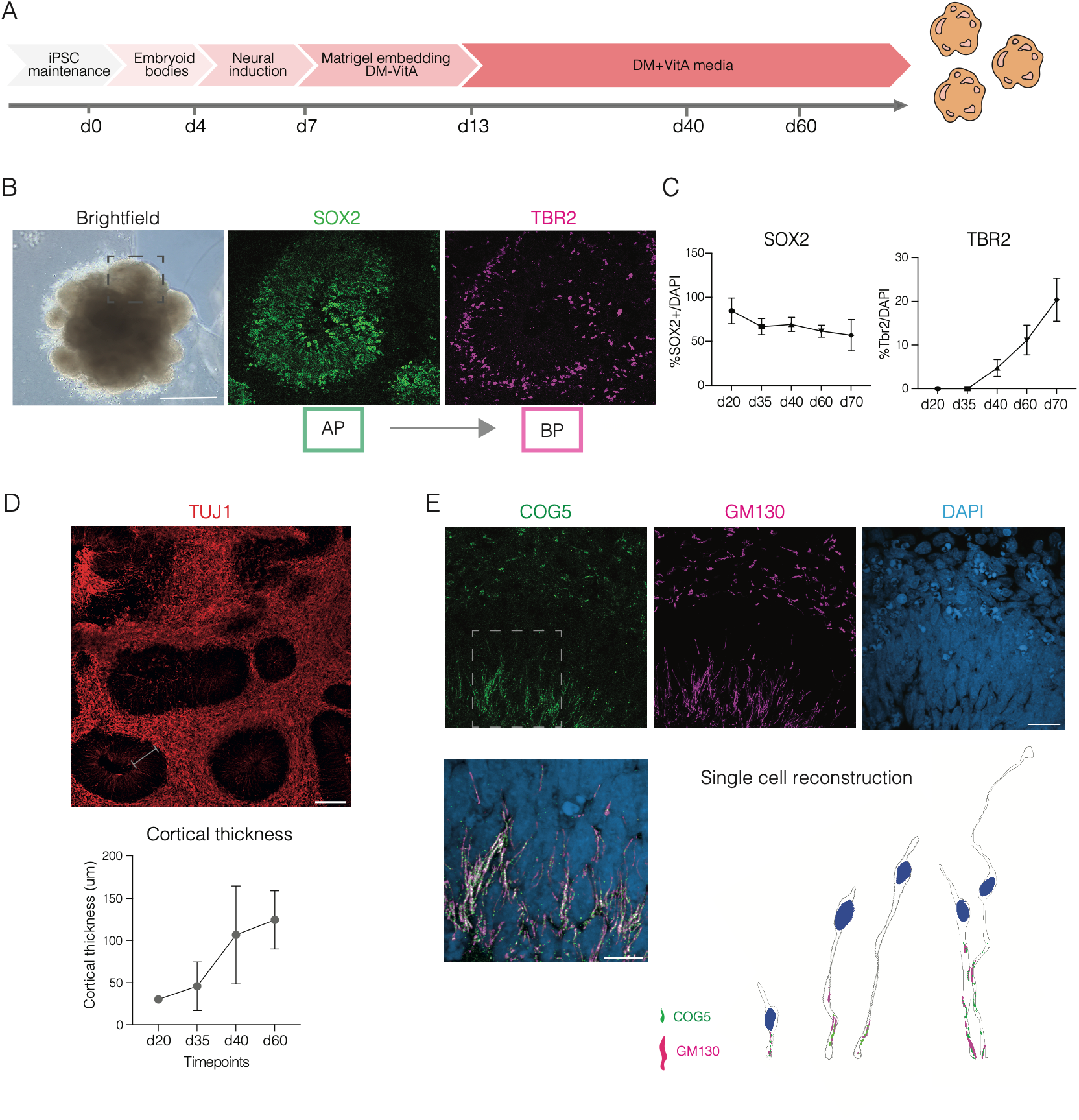
Characterization of human brain organoids and COG5 localization. **(A)** Experimental workflow for the generation of human brain organoids from induced pluripotent stem cells (iPSCs) following published protocol (see Material and Methods), including embryoid body formation, neural induction, Matrigel embedding, and differentiation stages. **(B)** Brightfield images at day 14. Scale bar 500 μm. Representative immunofluorescence images of organoid sections stained for SOX2 and TBR2, identifying apical progenitors (APs) and basal progenitors (BPs), at day 40. Scale bar 100 μm. **(C)** Quantification of percentage of SOX2⁺ and TBR2⁺ progenitors in organoids across developmental stages (days 20, 35, 40, 60, and 70). N=3 independent biological replicates. N=3 organoids. **(D)** Representative immunofluorescence images of organoid stained for TUJ1 at day 40. Scale bar 100 μm. Quantification of cortical thickness (organoids’ rosette width from the ventricular surface to the signal of TUJ1) across developmental stages (days 20, 35, 40, and 60). N=3 independent biological replicates. N=3 organoids. **(E)** Representative immunofluorescence images of organoids stained for the Golgi marker GM130, COG5, and DAPI. COG5 displays a punctate distribution colocalizing with Golgi compartments in apical progenitors. Scale bar 20 and 10μm. Representative reconstructions of single APs showing the distribution of COG5 and GM130 along the apical process.

**Supplementary Figure 5.**
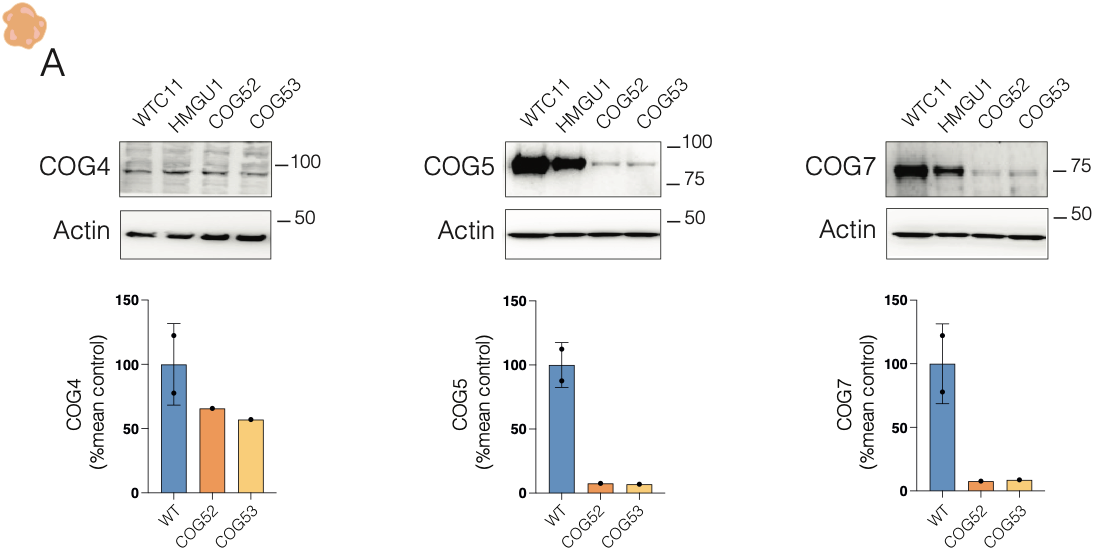
COG complex subunits proteins levels in organoids. **(A)** Western blot analysis and quantification of COG4, COG5, COG7 subunits expression levels in control and COG5-deficient organoids. N=1 independent organoid batch and n=3 total organoids per each condition.

**Supplementary Figure 6.**
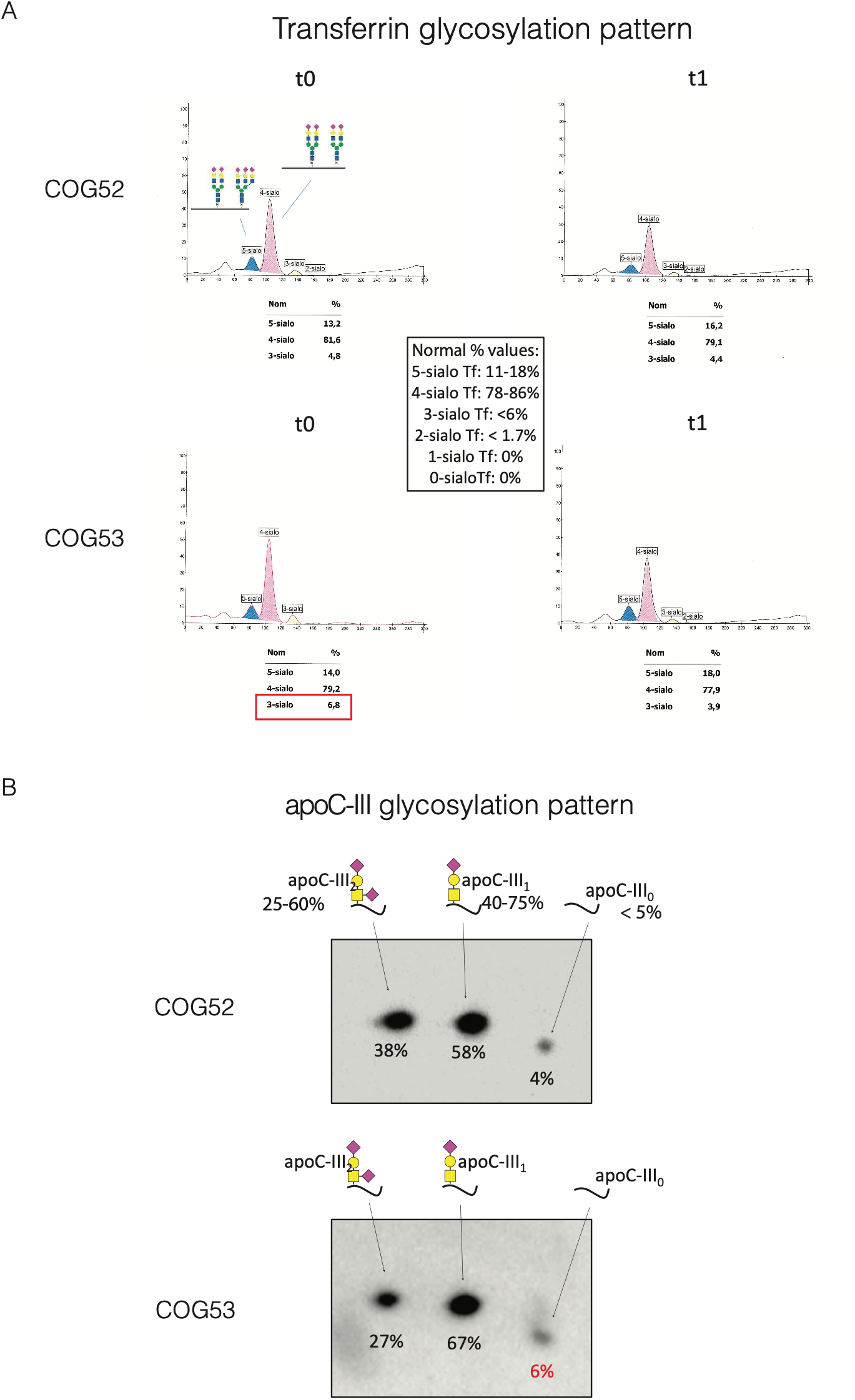
Plasma glycoprotein glycosylation profiles in COG5-CDG patients. **(A)** Transferrin glycosylation analysis performed on plasma samples from COG52 and COG53 at two independent time points (four months between the two analyses). Relative abundance of transferrin glycoforms is shown together with reference values. Both patients display largely preserved N-linked glycosylation profiles, with no major abnormalities detected by standard clinical transferrin analysis. **(B)** ApoC-III glycosylation analysis showing the relative abundance of apoC-III glycoforms in plasma samples from COG52 and COG53. O-linked glycosylation profiles are largely preserved, with only minor deviations from reference values.

**Supplementary Figure 7.**
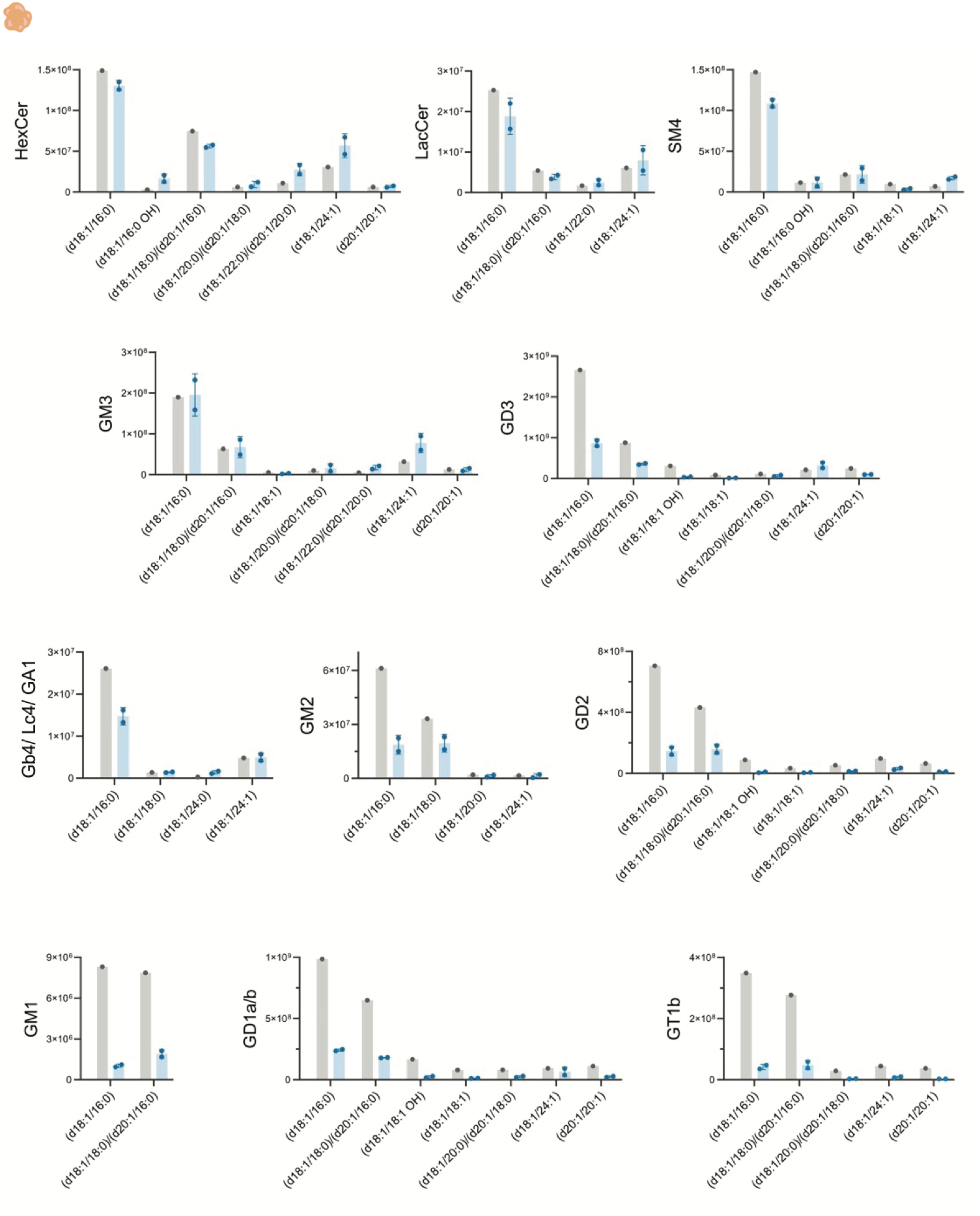
Glycosphingolipid (GSL) analysis in COG5-CDG brain organoids. GSL relative abundances (MS intensities) in control (WTC11; n=1) and COG5-deficient (n=2) organoids dissociated at day 40 for the predominant ceramide backbones. Each sample was generated by pooling 8 individual organoids together. Intensities are normalized to the internal standard and protein content per sample. Data are represented as the mean ± SD. Gangliosides GD3, GD2, GD1a/b, and GT1b represent the predominantly detected GSLs. GD1a and GD1b are structural isomers with identical exact masses and are not chromatographically resolved, so they were measured as a combined signal by MS (referred to as GD1a/b). Gb4, Lc4 and GA1 have the same mass-to-charge (m/z) ratio and are measured together. HexCer: Hexosylceramide (Glucosylceramide + Galactosylceramide), LacCer: Lactosylceramide; SM4: sulfatide.

**Supplementary Figure 8.**
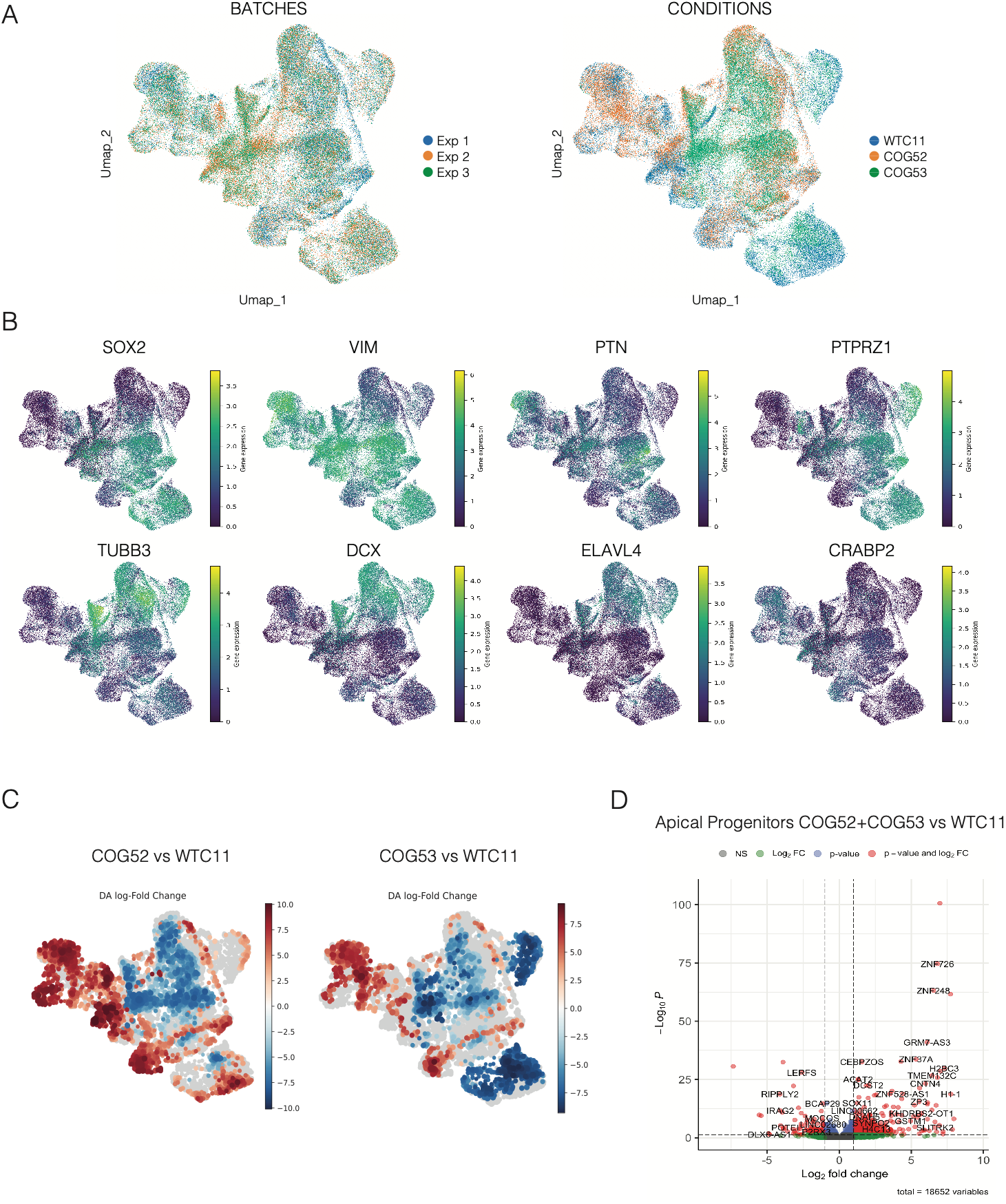
Quality check analyses and differential gene expression analysis of control and COG5-deficient organoids. **(A)** UMAP representation of the integrated single-cell RNA sequencing dataset for experimental batch (left) and genotype (right), showing successful integration and distribution of samples across identified cell populations. **(B)** Feature UMAP plots showing expression of representative marker genes used for cluster annotation, including markers of progenitors, neurons, and extracellular matrix-associated cell states. **(C)** UMAP representation of the differential abundance (DA) analysis comparing COG52 and COG53 organoids with wild-type controls. Positive (in red) and negative (in blue) DA log-fold change values identify cell populations enriched or depleted in COG5-deficient organoids relative to controls. **(D)** Enhanced volcano plots showing differentially expressed genes in apical progenitors comparing COG5-deficient organoids with wild-type controls. Representative random significantly upregulated and downregulated genes are labelled.

**Supplementary Figure 9.**
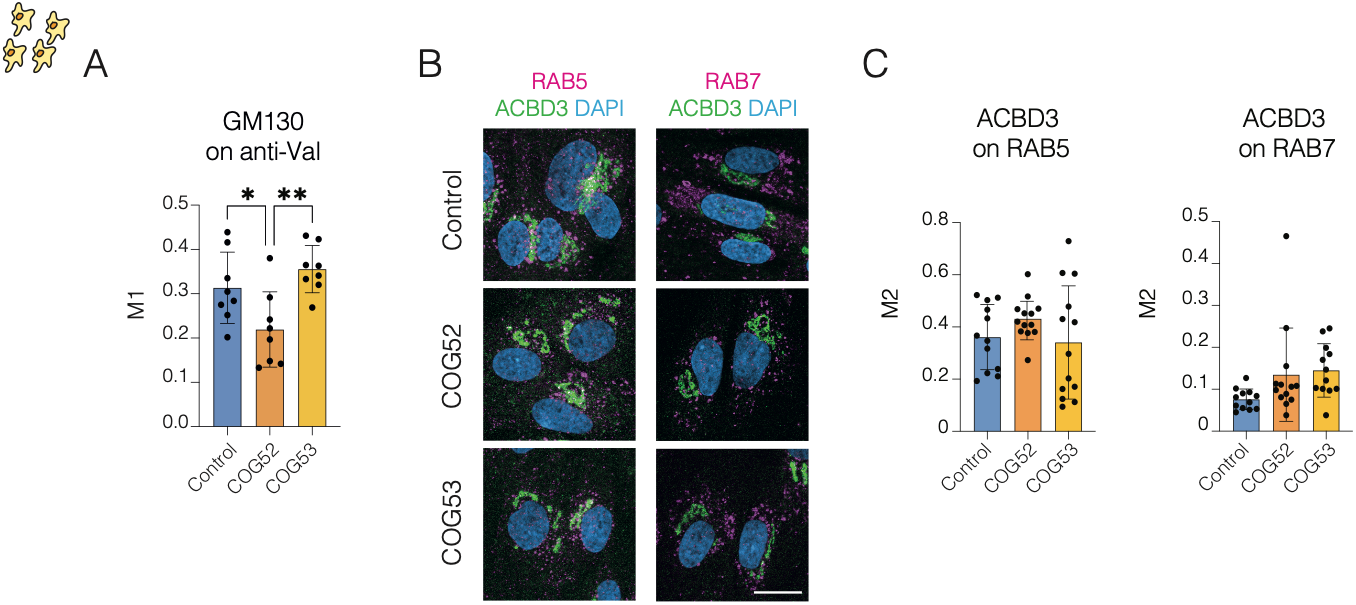
ACBD3 endosomal localization. **(A)** Colocalization quantified as M1 value from JacoP plugin in Fiji. N=1 independent experiment and n=3-10 total images quantified per each condition. One-way ANOVA, Tukey post test, *p<0,05, **p<0,01. **(B)** Representative images of ACBD3 localization relative to early and late endosomes (Rab5 and Rab7) in control and COG5-deficient fibroblasts. Scale bar 20μm. **(C)** Colocalization quantified as M1 value from JacoP plugin in Fiji. N=2 independent experiments and n=12-18 total images quantified per each condition. One-way ANOVA, Tukey post test.

## CONTRIBUTIONS

M.P. and E.T. conceived the experiments. E.T. supervised the project. M.P. performed cell culture, organoids culture, organoids manipulation, microscopy, image analysis and quantifications. A.J.P., B.G. and P.W. performed MS and glycolipidomics. E.R. performed fractionations, immunoblot and western blot. E.R and S.A. performed immunofluorescence on patients’ fibroblasts. V.R. and G.V. performed organoids culture. M.G., C.C., S.B. R.T. analyzed the scRNAseq data. F.F. kept in culture patients’ fibroblasts and performed immunofluorescence upon BFA treatment. M.W., I.T.A., O.P., F. A., M.M., S.V.B., A.B. were in charge of all clinical work and patients’ fibroblasts collection. M.P. and E.T. wrote and edited the manuscript. M.G, and V.R. co-wrote the final version of the manuscript,

## ACKNOWLEDGEMENTS

We are grateful to the services and facilities of HT for the outstanding support provided, notably, N. Maghelli and the team of the National Facility (NF) of Light Imaging, G. Faga’ and the team of Genome Editing and Disease Modeling Facility. We thank Thomas Vaccari and Elisabetta Argenzio for the critical reading of the manuscript and all members of the Taverna lab for helpful discussions.

## FUNDINGS

M.P. is a PhD student within the European School of Molecular Medicine (SEMM) at the University of Milan.

## Notes

### Competing Interest Statement

The authors have declared no competing interest.

